# Importin α/β1 dependent nuclear import of Black Sea Bass Polyomavirus Large Tumor Antigen is mediated by a classical NLS located downstream of the SF3 helicase domain

**DOI:** 10.1101/2025.05.20.655083

**Authors:** Mikayla Hoad, Silvia Pavan, Sepehr Nematollahzadeh, Ole Tietz, Jospeh Reeman, Jade K. Forwood, Gualtiero Alvisi

## Abstract

Polyomaviruses (PyVs) are small dsDNA viruses that replicate in the host cell nucleus, primarily relying on the viral encoded large tumor antigen (LTA). Aided by recent advances in molecular biology techniques, the list of known PyVs is rapidly growing, revealing unexpected broad sequence, and host heterogenicity. Given their dependence on nuclear localization, large tumor antigens represent an attractive model for studying the nuclear transport process. A comprehensive analysis of the evolution of classical nuclear localization signals (cNLSs) within LTAs encoded by PyVs infecting mammals highlighted strong positional conservation of cNLSs between the LXCXE motif and the origin-binding domain (OBD). Here we extend such analysis to PyVs infecting non-mammalian hosts. We combined biochemical, structural and functional assays to demonstrate that Black Sea Bass (BSB) PyV-LTA is transported into the nucleus by the Importin (IMP)α/β1 heterodimer thanks to the recognition of a bipartite cNLS located downstream of the SF3 helicase domain, rather than between the LXCXE motif and the OBD. Such cNLS binds with high affinity to several IMPα isoforms by simultaneously interacting with the minor and major binding sites. Substitution of NLS key basic residues abrogating binding to IMPα, or co-expression with the well characterized IMPα/β1 inhibitor Bimax2 suppressed nuclear localization. Intriguingly a cNLS could be identified in a similar position in LTAs from other PyVs infecting ray-finned fishes, but not cartilaginous fishes, birds or scorpions, where cNLSs were predicted elsewhere. Our study suggests that LTAs from PyVs infecting different non-mammalian hosts might bear cNLS in distinctive positions, possibly reflecting processes of virus-host adaptation.

## INTRODUCTION

Since the identification of the Murine polyomavirus (MPyV) in 1960 as a filterable agent inducing many different tumors in experimental animals (1), more than one hundred additional PyVs have been discovered, including John Cunningham (JC)PyV, BKPyV and Merkel Cell (MC)PyV, which are well-characterized human pathogens (2). Once believed to exclusively infect mammals, PyV sequences have been subsequently retrieved by a wide range of evolutionary distinct hosts, including birds, fishes and arachnids. The first fish PyV was identified from black sea bass (BSB) (3). This was quickly followed by the identification of PyVs from a giant guitarfish and the sharp-pined notothen (4). Subsequently, novel PyVs were identified from a Gilt-head bream (5) and an emerald notothen (6). Lastly, a PyV sequence could be retrieved from a thornback skate (7). Up to date, the 115 ICTV classified PyVs are assigned to 8 distinct genera, while a few remain still unclassified (8). PyVs are believed to have co-evolved with their host for over 500 million years, through complex mechanisms which likely involve intra-and inter-family recombination (9). PyVs appear to be highly species-specific, as highlighted by the lack of documentation of human-to-human transmission of simian virus 40 (SV40), which infected millions of US residents via contaminated antipolio vaccine stocks (10). Therefore, PyVs are believed to have originated from at least one ancestral virus that infected the last common ancestor of arthropods and vertebrates, co-evolving with their hosts and diverging from one another at a faster rate than host speciation, through an intrahost divergence model, with occasional adaptation to new hosts (4). Accordingly, several of the 14 human (H)PyVs described are evolutionary distant and not monophyletic, being assigned to three different genera (2). For example, the JC and BKPyVs, the first HPyVs discovered, are more closely related to the resus macaque monkeys-infecting SV40 than to the other HPyVs. All PyVs are small dsDNA viruses, which are believed to replicate into the host cell nucleus thanks the concerted action of cellular and viral proteins. Among the latter, the large tumor antigen (LTA) is a multifunctional viral product which has been mainly characterized for SV40 (11). SV40-LTA contain several well-characterized domains, which appear loosely conserved amongst HPyVs (11). At the N-terminus, a J domain, containing the HPDKGG hexapeptide motif, is essential for interactions with host chaperone proteins such as Hsp70, crucial for viral replication and the transformation of host cells (12). Adjacent is the LXCXE motif, which enables binding to retinoblastoma (Rb) family proteins, disrupting cell cycle regulation and promoting viral replication (13). Continuing downstream is the origin-binding domain (OBD) which specifically binds to the origin of replication of SV40 DNA, initiating viral DNA replication (14). Lastly, at the C-terminus, the Tag-D1 type Zn-binding domain - responsible primarily for the formation of T antigen hexamers (15), and the SF3 helicase domain, possessing the ATPase/helicase properties needed for unwinding the viral DNA during replication (16). Despite being mainly known for its transforming properties in cell culture and its ability to induce tumors when exogenously expressed in nonpermissive immunodeficient animal species (17), SV40-LTA has been extraordinarily important for the study of nucleocytoplasmic transport. The first nuclear localization signal (NLS) was identified between the LXCXE motif and the OBD of SV40-LTA, as a sequence both essential for nuclear targeting of LTA and sufficient to mediate nuclear localization to otherwise cytoplasmic reporter proteins (18–20). Today the SV40-LTA NLS is considered the prototype of the so-called classical(c)NLSs, able to confer nuclear localization by being recognized by the importin (IMP)α/β1 heterodimer, whereby IMPα recognizes the NLS-bearing cargoes and IMPβ1 mediates docking and translocation across the nuclear pore complex (NPC, see (21) for a review). cNLS can be either monopartite - formed by a single stretch of basic amino acids interacting with the IMPα major binding site and exemplified by the SV40-LTA NLS (22), or bipartite-comprising two basic stretches of amino acids, interacting with IMPα minor and major binding sites simultaneously, and exemplified by the nucleoplasmin bipartite NLS (23). The study of nuclear transport mechanisms of LTAs from PyVs infecting mammals allowed to propose a model for the evolution of bipartite NLSs by duplication of a monopartite NLS (24). Indeed, cNLSs have been identified between the LXCXE motif and the OBD in LTAs from all classified PyVs infecting mammals, whereby the SV40 cNLS remains highly conserved. The main difference lies in NLS duplication events which resulted in c. 50% of LTAs bearing a bipartite NLSs. However, no functional cNLS could be bioinformatically predicted in such position for LTAs from PyVs infecting non-mammalian hosts, providing a unique opportunity to further study NLS evolution (24). Here we combine structural and biochemical assay with confocal laser scanning microscopy (CLSM) to address the issue of NLS evolution between highly divergent LTAs and thoroughly characterize a bipartite cNLS in BSB PyV-LTA, which is conserved in other PyVs infecting ray-finned fishes. Therefore, despite although the existence of c. 20 orthologues of IMPβ1, capable of recognizing specific NLSs (25), PyVs converge on the IMPα/β1 heterodimer to ensure nuclear transport of LTAs. A bioinformatics analyses identified a potential cNLS in all of LTAs regardless of their divergence. However, their position is highly variable and distinctive of the host species infected, with LTAs from PyVs infecting ray-finned fishes, cartilaginous fishes, birds and scorpions bearing cNLS in different locations.

## Materials and Methods

### Plasmids

Open reading frames encoding for the aminoacidic sequence of LTA from SV40 (UniProt code: # P03070) or BSB PyV (UniProt code: A0A0A1C4T1) were optimized for human expression with Geneoptimizer software (Thermo Fisher Scientific, Monza, Italy). Subsequently, sequences were synthesized and cloned in plasmid pEGFP-N1 by GenScript (Singapore) to obtain plasmids pEGFP-N1-SV40-LTA and pEGFP-N1-BSB PyV-LTA. Plasmids pEGFP-N1-BSB PyV-LTA;mNLSn, pEGFP-N1-BSB PyV-LTA;mNLSc and pEGFP-N1-BSB PyV-LTA;mNLSnc, containing Alanine substitutions within either BSB PyV-LTA N-terminal, C-terminal or both basic stretches forming its putative bipartite NLS were obtained by site-specific mutagenesis of pEGFP-N1-BSB PyV-LTA by GenScript (Singapore). Plasmid mcherry-Bimax2, encoding a competitive inhibitor of the IMPα/β1 nuclear import pathway (26, 27), was kindly gifted from Yoshihiro Yoneda and Masahiro Oka (Osaka, Japan). Plasmids encoding IMPs used for electromobility shift assays (EMSAs) and fluorescence polarization (FP) assays include human α1 (hIMPα1ΔIBB; His tag, TEV site, UniProt code: P52292), α3 (hIMPα3 ΔIBB; His tag, TEV site, UniProt code: O00629), α5 (hIMPα5ΔIBB; His tag, TEV site, UniProt code: P52294), α7 (hIMPα7ΔIBB; His tag, TEV site, UniProt code: O60684), mouse α2 (mIMPα2ΔIBB; His tag, no TEV site, UniProt code: P52293), human β2 (hIMPβ2; His tag, TEV site, UniProt code: Q92973), and human β3 (hIMPβ3; His tag, TEV site, UniProt code: O00410) in pET30a (28), and human β1 (hIMPβ1; His tag, TEV site, Uniprot: Q14974) in pMCSG21 (29). A list of all plasmids used is available in Supplementary Table I.

### Peptides

Synthetic peptides including an Ahx-FITC modification of the N-terminus were synthesized at Macquarie University, Sydney, Australia, using standard Fluorenyl methoxycarbonyl (Fmoc)-solid-phase peptide synthesis on a CEM Liberty Blue™ Peptide Synthesiser (CEM, USA). Briefly, rink amide resin was pre-swelled in 50/50 dimethylformamide (DMF) and dichloromethane (DCM) for 1 hour. Amino acids were dissolved in DMF at a concentration of 0.2 M before being transferred to the synthesiser. Peptides were synthesized using sequential amid coupling from C to N-terminus for 3 minutes at 90°C, using five equivalents of amino acid with 10 equivalents of activator (0.5M DIC (N,NII-Diisopropylcarbodiimide) in DMF) and 5 equivalents of activator base (0.5 M Oxyma (Ethyl cyanohydroxyiminoacetate), 0.05 M DIPEA (N,N-Diisopropylethylamine) in DMF, followed by Fmoc deprotection in 20% piperidine in DMF for 2 minutes at 90°C and resin wash in DMF. Double couplings were performed for arginine residues to ensure complete coupling. Following final Fmoc deprotection of N-terminal amino hexanoic acid (Ahx), the resin was removed from the synthesiser, transferred to a syringe fitted with a propylene filter, washed, and FITC coupling was performed. FITC is coupled using 3 equivalents of FITC and 6 equivalents of DIPEA in DMF overnight. Peptides were washed with DMF, DCM, and methanol before cleavage. Peptides were cleaved from resin using cleavage cocktail of 92.5% TFA (trifluoroacetic acid), 2.5% TIPS (triisopropylsilane), 2.5% thioanisole and 2.5% H_2_O for 3-6 hours at room temperature, precipitated in ice cold diethyl ether, dissolved in H_2_O, freeze dried, and purified using a Shimadzu LC-20AD High-performance liquid chromatography (HPLC, Shimadzu, Japan). Mass spectra were obtained on a Shimadzu LCMS-8050 system (Shimadzu, Japan) in positive electron spray [ESI+] mode, fitted with a Polaris 3 C18-A 150 x 4.6 mm column (Agilent Technologies, USA). Peptides designed included BSB PyV-LTA;NLS and the three mutated versions with Alanine in place of basic residues. These mutation peptides are designated; BSB PyV-LTA;mNLSn, BSB PyV-LTA;mNLc, and BSB PyV-LTA;mNLSnc. A list of peptides used is available in Supplementary Table II. Analytical data from the production of the peptides can be found in the supplementary figures: BSB PyV-LTA;NLS (Supplementary Figure S1), BSB PyV-LTA;mNLSn (Supplementary Figure S2), BSB PyV-LTA;mNLSc (Supplementary Figure S3), and BSB PyV-LTA;mNLSnc (Supplementary Figure S4).

### Purification of recombinant proteins

Plasmids encoding for recombinant IMPα and IMPβ isoformswere transformed via heat shock method (30) into competent BL21(*DE3*)pLysS *E. coli* cells (#EC0114, Thermo Fisher Scientific, Monza, Italy) and recombinantly expressed via large scale auto-induction (31) expression method for 30 – 36 hrs at room temperature (25°C). After induction, expression cultures were centrifuged at 6000 rpm for 30 minutes at 16°C, and the resulting bacterial pellets were resuspended in HIS buffer A (50 mM phosphate buffer, 300 mM NaCl, and 20 mM imidazole, pH 8.0). Cells were lysed by two freeze-thaw cycles, followed by the addition of lysozyme (50 mg/mL; #L6876, Sigma-Aldrich, St. Louis, MI, USA) and DNase (20 mg/mL; #DN25, Sigma-Aldrich, St. Louis, MI, USA), and incubated at room temperature for 1 hour on a roller. Supernatants containing soluble proteins were collected by centrifugation at 12000 rpm for 45 minutes at 4°C. The extracts were then filtered through 0.45 μm low protein affinity filters and using an AKTA purifier FPLC system (Cytiva, Washington D.C., USA) injected onto a 5 mL HisTrap HP column (#17524801, Cytiva, Washington D.C., USA) that had been pre-equilibrated with His buffer A. Followed by 20 column volumes washes with His buffer A, recombinant proteins were eluted using a gradually increasing gradient of imidazole (ranging from 20 mM to 500 mM) (#288-32-4, ChemSupply, Gillman, SA, Australia). The eluted protein fractions were combined and loaded onto a pre-equilibrated HiLoad 26/60 Superdex 200 column (#28989336, Cytiva, Washington D.C., USA) in GST buffer A (50 mM Tris and 125 mM NaCl) for further purification using size-exclusion chromatography. The fractions corresponding to the eluted volumes at the respective protein sizes were collected, and the samples were concentrated using an Amicon MWCO 10 kDa filter (#UFC9010, Merck Millipore, Burlington, MA, USA). Before experimental use, the purity of the samples was assessed by performing SDS-PAGE at 165 V for 30 minutes on a 4–12% Bis-Tris Plus gel (#NW04120BOX, Thermo Fisher Scientific, Waltham, MA, USA).

### Electrophoretic mobility shift assays (EMSAs)

Qualitative binding analysis using native gel was performed using EMSAs. IMPs were electrophoretically separated in the presence or in absence of BSBPyV LTA FITC-NLS peptides to determine protein:peptide binding interactions. A 1.5% agarose gel in TB buffer (0.45 mM tris, 0.45mM boric acid, pH ~8.5) was loaded with recombinant proteins (20 μM) and/or BSB PyV-LTA;NLS peptides (10 μM) and run for 2 hours at 75 V. Gel was imaged twice, once with a UV filter to detect fluorescence of peptides and a second time with visible light to detect Coomassie stained protein bands.

### Fluorescence polarization assays (FPs)

FITC-tagged NLS peptides (10 nM) were incubated with recombinant IMPs in two-fold serial dilutions, starting from 20 μM, across 23 wells to a final volume of 200 μL per well in GST buffer A (50 mM Tris, 125 mM NaCl), with the last well not containing any IMPs, essentially as described previously (24, 32–35). FP measurements were performed using a CLARIOstar Plus plate reader (BMG Labtech, Germany) in 96-well black Fluotrac microplates (#655076, Greiner Bio-One, Austria). Data were analyzed with GraphPad Prism (Prism 9, Version 9.3.1), and a binding curve was fitted to the one-site-specific binding least square fit function to determine the dissociation constant (Kd) and maximum binding (Bmax).

### Crystallography

Protein crystals of IMPα2 were produced using the hanging-drop vapour diffusion method (36) in a range of known conditions (0.50 – 0.85 M sodium citrate, 0.10 M HEPES pH 6.5/7.0/7.5, and 0.01 M DTT) at a concentration of 18 mg/mL. Grown IMPα2 crystals were soaked with BSB PyV-LTA;NLS peptide three hours before harvesting. Soaked crystals were cryo-protected in a reservoir solution containing 25% glycerol for 10 seconds before flash freezing in liquid nitrogen. Crystals were diffracted at the Australian Synchrotron on the MX2 (37) beamline. Diffraction data was processed using DIALS (38), merged and scaled through Aimless (39) phased using molecular replacement with model 6BVT (40) from the protein data bank (PDB) (41) and finalized with iterative cycles of refinement through Phenix (42) and structural modelling in Coot (43). The final structure was validated and deposited on the PDB with the ID code 9NIB. All crystallization, data collection and refinement statistics are listed in Supplementary Table III. Interface interactions between BSB PyV-LTA;NLS and IMPα used in the structure figures were calculated using the PDBsum analysis program (44). A full list of calculated binding interactions is listed in Supplementary Table IV. The final structural model was solved to a resolution of 2.2Å and refined to a R_work_/R_free_ of 0.19/0.22. Final model was resolved with 424 residues of IMPα2, 12 residues of BSB PyV-LTA;NLS and 86 water molecules.

### Cells

HEK293A cells (#R70507, Thermo Fisher Scientific, Monza, Italy) were maintained in Dulbecco’s modified Eagle’s medium (DMEM) supplemented with 10% (v/v) fetal bovine serum (FBS), 50⍰U/mL penicillin, 50⍰U/mL streptomycin, and 2⍰mM1l-glutamine in a humidified incubator at 37°C in the presence of 5% CO_2_ and passaged when reached confluence (45).

### Transfections

HEK293A cells were seeded in a 24-well plate onto glass coverslips (5⍰× ⍰ 10^4^ cells/well). The following day, cells were transfected with appropriate expression plasmids (range 125–250⍰ng), using 1 μl Lipofectamine 2000 (#11668019, Thermo Fisher Scientific, Monza, Italy), and 100 μl OptiMEM per well. Cells were incubated for 24 in a humidified incubator at 37°C in the presence of 5% CO_2_ using DMEM devoid of antibiotics until being processed for CLSM (46, 47).

### CLSM and Image Analysis

Cells were incubated with DRAQ5 (#62251, Thermo Fisher Scientific, Monza, Italy) at a dilution of 1:5000 in DMEM without phenol red for 30⍰min in a humidified incubator at 37°C in the presence of 5% CO_2_. Subsequently, cells were washed twice with PHEM 1× solution (60⍰mM PIPES, 25⍰mM HEPES, 10⍰mM EGTA, and 4⍰mM MgSO_4_) and fixed with 4% paraformaldehyde (#I28800, Thermo Fisher Scientific, Monza, Italy) for 10 min at room temperature. After three washes with PBS 1×, coverslips were mounted on glass slides using Fluoromount G (#00-4958-02, Thermo Fisher Scientific, Monza, Italy), as previously (48). For each condition, z stacks of several randomly chosen fields with a total thickness of 2 μm and a step of 0.4 μm were acquired with a Nikon A1 confocal laser scanning microscope (Nikon, Tokyo, Japan) equipped with a 60× oil immersion objective, using the software NIS elements (Nikon, Tokyo, Japan). Maximum intensity projections were generated and used to measure the levels of nuclear accumulation at the single cell level with the FiJi public domain software (https://doi.org/10.1038/nmeth.2019). To this end, single-cell measurements were taken for nuclear (Fn), and cytoplasmic (Fc) fluorescence. DRAQ5 was used to define nuclear, whereas a small area close to DRAQ5 was used to define a cytosolic mask. The fluorescence attributed to autofluorescence/background (Fb) was subtracted from the measurements to calculate the Fn/c ratio according to the formula Fn/c⍰= ⍰(Fn⍰-⍰ Fb)/(Fc⍰-⍰ Fb) as previously described (49, 50). Cells with oversaturated signals were excluded from the analysis.

### Statistical analysis

Statistical analysis was performed using GraphPad Prism 10.1.0 software (GraphPad, San Diego, CA, USA) applying Student’s t-test, one-way ANOVA, or two-way ANOVA, as appropriate.

### Phylogenetic analysis

LTA sequences were retrieved from UniProt. Sequences were aligned using clustal Omega (51) and a phylogenetic tree weas constructed using MEGAX with the Neighbor Joining method (52).

### Bioinformatics

cNLS sequences were identified using the software cNLS mapper (53) with a cutoff score of 5 and including bipartite NLSs with a long linker (13-20 aa) within the full aminoacidic sequence. Functional domains on LTAs were retrieved from UniProt (54) and Prosite (55). In specific cases, domains were either identified with HHPred (56) or structurally predicted by AlphaFold3 (57) with visualization and model coloring done using UCSF ChimeraX program (version 1.6.1).

## Results

### Identification of functional domains and a putative cNLS wthin BSB PyV-LTA

We have recently demonstrated that LTAs from all known HPyVs are transported into the nucleus by the IMPα/β1 heterodimer after recognition of cNLSs located between the LXCXE motif and the OBD. Intriguingly, a bioinformatics analysis failed to predict a cNLS in a similar position in LTAs from PyVs infecting non-mammalian hosts (24). Among them, the LTA from BSB PyV, the first PyV infecting fishes identified, is only 27.5 % identical to SV40-LTA, and despite sequence and structural homology-based algorithms predicted the presence of the typical LTA domains, it lacks a HPDGKK motif, so it is likely missing a functional J domain (Figure 1AB). Additionally, the LXCXE motif, which is commonly placed between the J domain and the OBD, is located within the OBD (Figure 1). Interestingly, despite a functional cNLS could not be predicted within the short linker connecting the putative J and OBD domains, it bears a TPEK peptide reminiscent of the functional cNLS described in LTAs from PyVs infecting mammals (24). Such sequence does not match the consensus for IMPα binding (K-K/R-X-K/R) (49) and therefore it is not likely responsible for nuclear transport of BSB PyV-LTA. However, our analysis identified a potentially very active cNLS (660-IAEYKRRHNIGSDGLPNKRRRCLF-683) within the largely unstructured C-terminus of the protein, located downstream of the SF3 helicase domain (Figure 1AB). This suggests that during the evolution of highly divergent LTAs, the dependence on the classical IMPα/β1 dependent nuclear import pathway is preserved, but the position of the cNLS changed.

**Figure 1.**
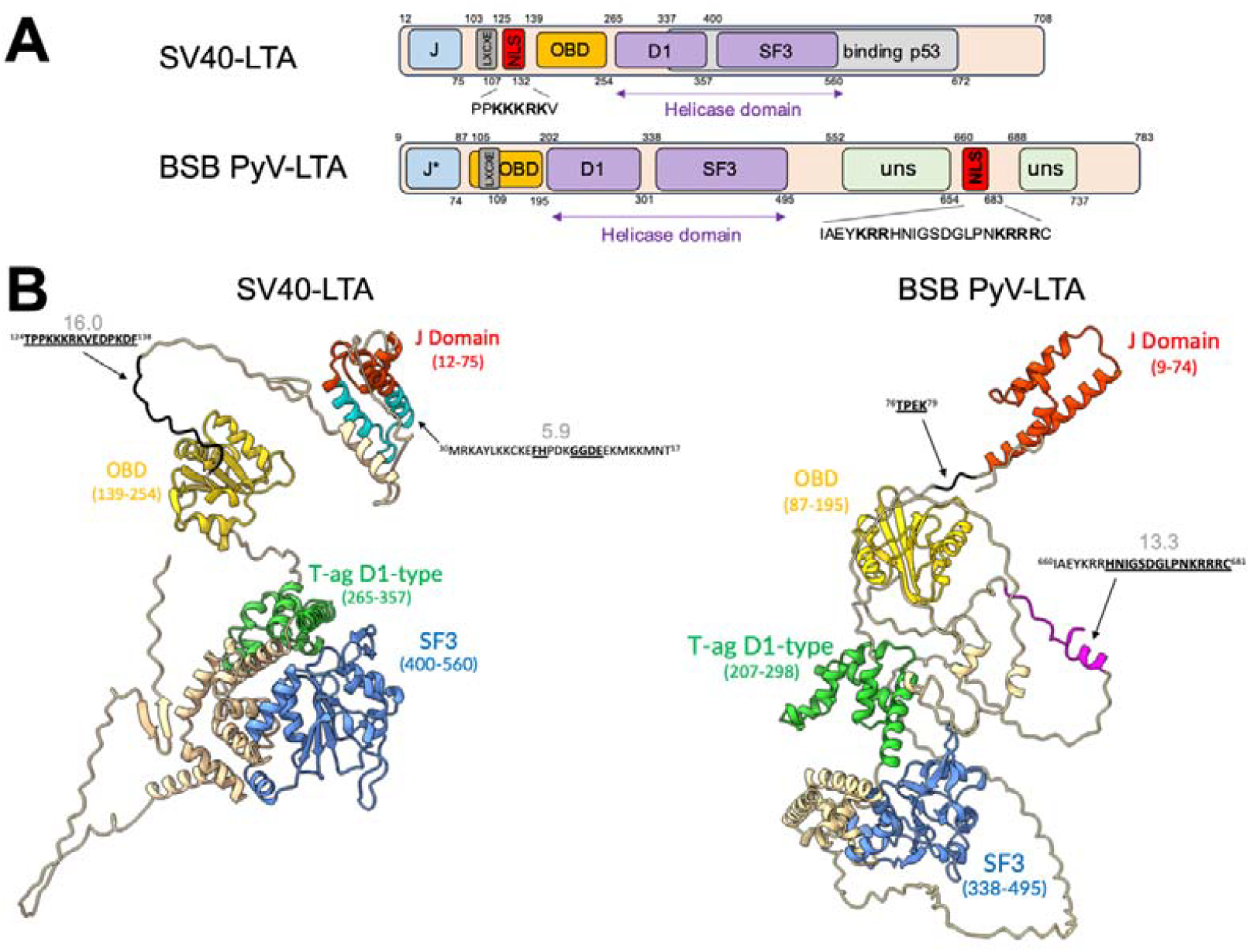
Functional domains and motifs predicted within BSB PyV-LTA. (A) Schematic representation of SV40 and BSB PyV-LTAs, along with predicted functional domains and motifs. The cNLS sequences are shown with the single letter amino acid code, with basic residues in boldface. An asterisk close to the predicted J domain of BSB PyV-LTA indicates the absence of the HPDGKK hexapeptide, suggesting lack of functionality. (B) 3D structures of SV40 (left) and BSB PyV-LTAs monomers predicted using AlphaFold3. The specific domains and motifs are colored according to the following scheme: J domain, red; origin binding domain (OBD), yellow; T-ag D1 type, green; helicase domain (SF3), blue. cNLSs have been indicated with arrows, and colored according to their position: within J domain, cyan; upstream of the OBD, black; downstream of the SF3 helicase domain, magenta. cNLS sequences are shown with residues in bold and underlined representing flexible regions of the structure whilst those not bold or underlined are identified to be part of the predicted secondary structures. The cNLS mapper score relative to each putative cNLS is indicated in gray.

### BSB PyV-LTA residues 660-683, located downstream of the SF3 helicase domain interact with IMPα isoforms with high affinity

To validate this hypothesis, we assessed the ability of BSB PyV-LTA;NLS to bind several NTRs, including IMPαΔIBB from each IMPα subfamily (SF1: IMPα1, and IMPα2, SF2: IMPα3, and SF3: IMPα5 and IMPα7) and IMPβ isoforms 1-3. To this end, a FITC-labelled BSB PyV-LTA;NLS peptide was subjected to native gel EMSA in the absence or presence of recombinantly purified IMPs. Importantly, the peptide co-migrated with several IMPα isoforms, although to different extents, indicating its potential as cNLS. Surprisingly, the peptide also co-migrated with IMPβ1, IMPβ2, and IMPβ3 (Figure 2A). These results suggest that BSB PyV-LTA residues 660-683 could be a functional NLS. However, their ability to directly bind IMPβ1, 2 and 3, in addition to all IMPα isoforms tested, implied the possibility that BSB PyV-LTA evolved to simultaneously exploit multiple nuclear import pathways, as recently described for several cellular (25) and viral proteins (34, 35). To verify this hypothesis, we measured the binding affinity between the same IMPs used in the EMSA and BSB PyV-LTA;NLS via FP assays (Figure 2B). Our results confirmed that BSB PyV-LTA;NLS binds to all tested IMPs, although with much higher affinity for IMPα7, IMPα5 and IMPα2 (Kd in the low nanomolar range) as compared to IMPβ1-3, for which an accurate K_D_ value could not be calculated due to the low levels of binding. Overall, our results indicate that although BSB-PyV LTA residues 660-683 can interact with several IMPαs and IMPβs, it binds to the former with much higher affinity, implying its potential as a *bona fidae* cNLS.

**Figure 2.**
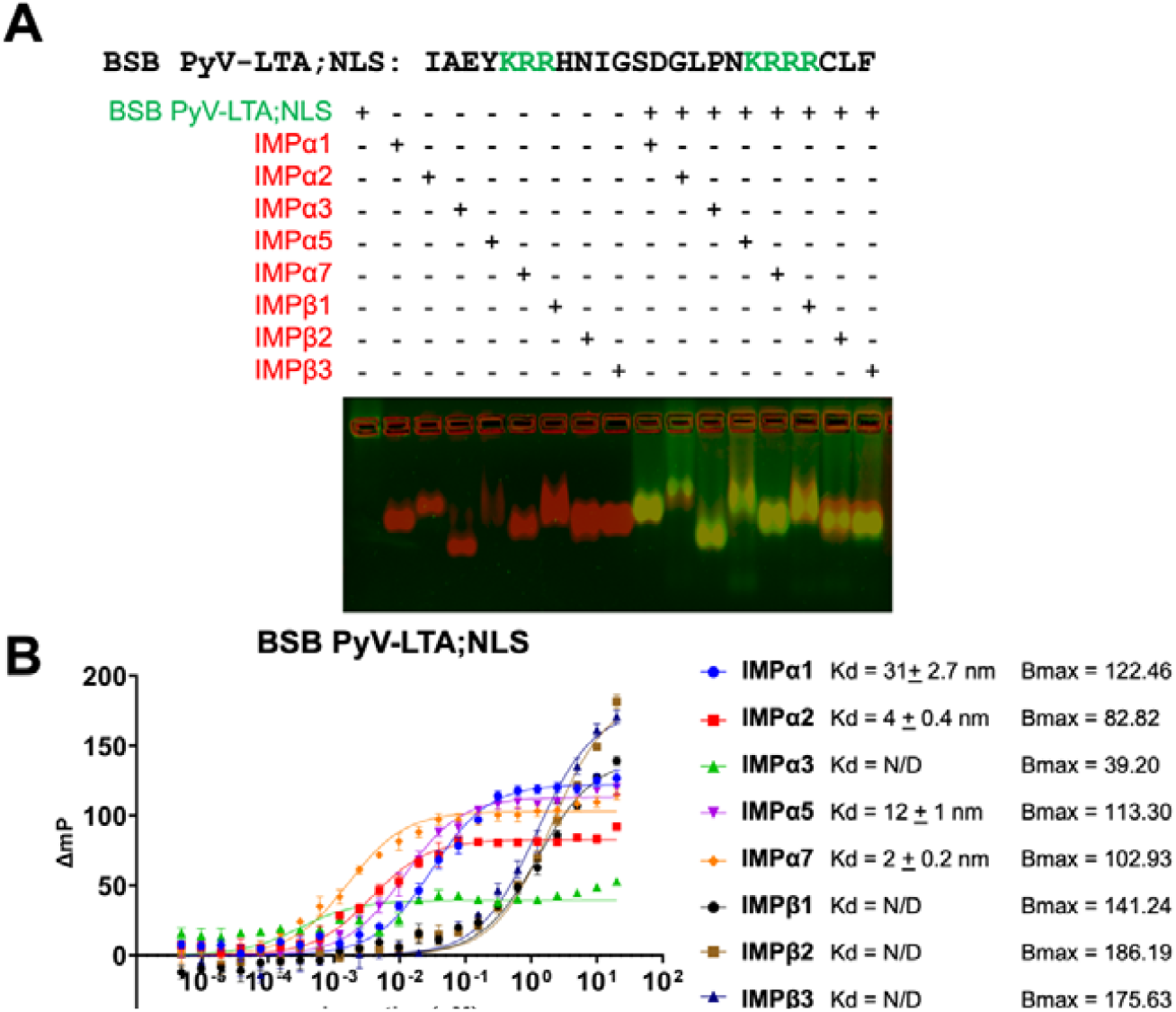
BSB PyV-LTA 660-683 interact with IMPα isoforms with high affinity. (A) EMSA for FITC labelled BSB PyV-LTA NLS peptide (10 μM) and recombinant IMPs (20 μM). After electrophoresis on 1.5% agarose gel, FITC labelled peptides were visualized with an UV filter (green), and proteins by Coomassie staining (red), with an image of both channels shown on bottom. (B) FP assays were performed by incubating the FITC labelled BSB PyV-LTA;NLS peptide (2 nM) with increasing concentrations of the indicated recombinant IMPs, followed by data acquisition and fitting to measure the dissociation constant (Kd) and maximum binding (Bmax) relative to each interaction. Data shown are mean ± standard error of the mean (SEM) relative to three independent experiments.

### BSB PyV-LTA is transported into the nucleus by IMPα/β1

The ability of BSB PyV-LTA;NLS to specifically bind with high affinity to IMPα isoforms suggests that the protein could be imported into the nucleus by the IMPα/β1 heterodimer, following a classical nuclear import pathway. We therefore investigated its subcellular localization in the presence or the absence of the IMPα/β1 competitive inhibitor Bimax2. To this end, BSB PyV-LTA-GFP was transiently expressed in a cellular context without any other viral protein, using SV40-LTA-GFP as a positive control (Figure 3A). As expected, SV40-LTA-GFP strongly accumulated in the cell nucleus (Figure 3B) of 100% of analyzed cells (Figure 3C) and an average Fn/c of 39.8 (Figure 3D). BSB PyV-LTA-GFP similarly accumulated in the nucleus in 100% of transfected cells, with a Fn/c of 7.8 (Figure 3B-D). Therefore, nuclear localization is conserved amongst highly divergent LTA infecting primates (SV40) and ray finned fishes (BSB). As expected, Bimax2 strongly inhibited nuclear localization of SV40-LTA-GFP (Figure 3B), resulting in a mainly cytoplasmic protein in 100% of co-transfected cells (Figure 3C), and an average Fn/c of 0.4 (Figure 3D). Importantly, a similar inhibition was be observed for BSB-PyV-LTA, which was retained in the cytosol of 100% of co-transfected cells with an average Fn/c of 0.3 (Figure 3BD).

**Figure 3.**
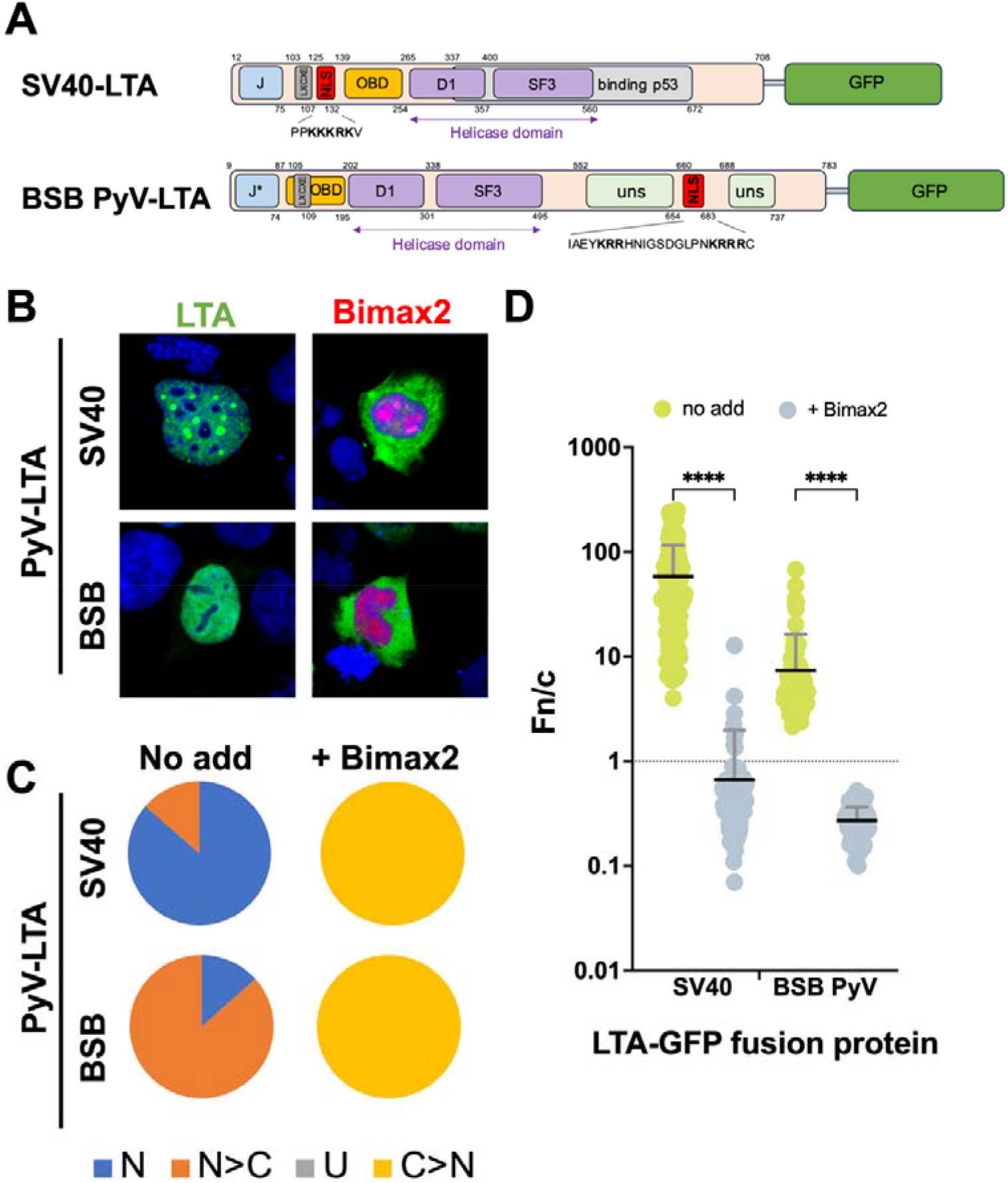
BSB PyV-LTA accumulates in the nucleus in the absence of any other viral protein in an IMPα/β1 dependent fashion. (A) Schematic representation of the GFP fusion proteins expressed. LXCXE: LXCXE motif; NLS: nuclear localization signal; OBD: origin-binding domain; D1: tag D1 type domain; SF3: SF3 helicase domain; uns: unstructured. (B) HEK293A cells were transfected with plasmids encoding the indicated LTA-GFP-fusions in the absence (*left* panels, LTA) or presence (*right* panels, Bimax2) of a plasmid encoding for mcherry-Bimax2. At 24 h p.t., cells were processed for CLSM and subcellular localization of GFP-fusion proteins was quantitatively analyzed by CLSM. The DRAQ5 (633 nm excitation laser line; blue), GFP (488 nm excitation laser line; green) and mcherry (561 nm excitation laser line; red) channels are shown in a merged image. (C) Micrographs, such as those shown in (A), were quantitively analyzed to calculate the Fn/c relative to individual cells (n > 90). Data shown are the percentage of cells displaying the indicated phenotype. N: Fn/c > 10; N>C: 2< Fn/c < 10; U: 1< Fn/c < 2; C: Fn/c < 1. (D) The mean ± standard deviation of the mean (SD) relative to each GFP-fusion, including the results of the student’s t-test for significance between expression in the absence (green circles) or the presence (purple circles) of mcherry-Bimax2. ****: p < 0.0005, *** p: < 0.005. The dotted horizontal line corresponds to Fn/c = 1, representing a ubiquitous distribution between nucleus and cytoplasm.

Therefore, although LTAs from BSB and mammals infecting PyVs possess a NLSs located in different domains, their nuclear import similarly depends on the IMPα/β1 heterodimer.

### Structural analysis of the BSB PyV-LTA;NLS:mIMPα2 complex reveals a bipartite mechanism of interaction

BSB PyV-LTA possesses between residues 660-683 two closely located stretches of basic amino acids which bind to IMPα isoforms high affinity, and it is translocated into the nucleus by the IMPα/β1 heterodimer. These observations strongly suggests that BSB PyV-LTA residues 660-683 represent a classical bipartite NLS simultaneously binding to both IMPα minor and major NLS binding sites. To investigate this possibility and gain structural insights regarding the interaction between BSB PyV-LTA and IMPα, we solved the crystal structure of BSB PyV-LTA residues 660-683 in complex with mIMPα2ΔIBB, which is normally used for such a purpose due to its ease of crystallization (Figure 4A and Supplementary Table III). The complex was solved with a 2.2 Å resolution and revealed BSB PyV-LTA residues 664-666 at the IMPα2 minor site, and residues 672-688 at the major binding site (Figure 4B and Supplementary Table IV), reminiscent of bipartite NLSs already described for several other LTA from mammalian-infecting HPyVs (24). In particular, the typical salt bridge between NLS side chain R665 and IMPα2 minor site GLU396 as well as between NLS side chain K675 and IMPα2 major site ASP192 were detected in the minor site P2’ pocket and the major site P2 pocket, respectively, whereas several other NLS side chains engaged in hydrophobic interactions (Figure 4B and Supplementary Table IV).

**Figure 4.**
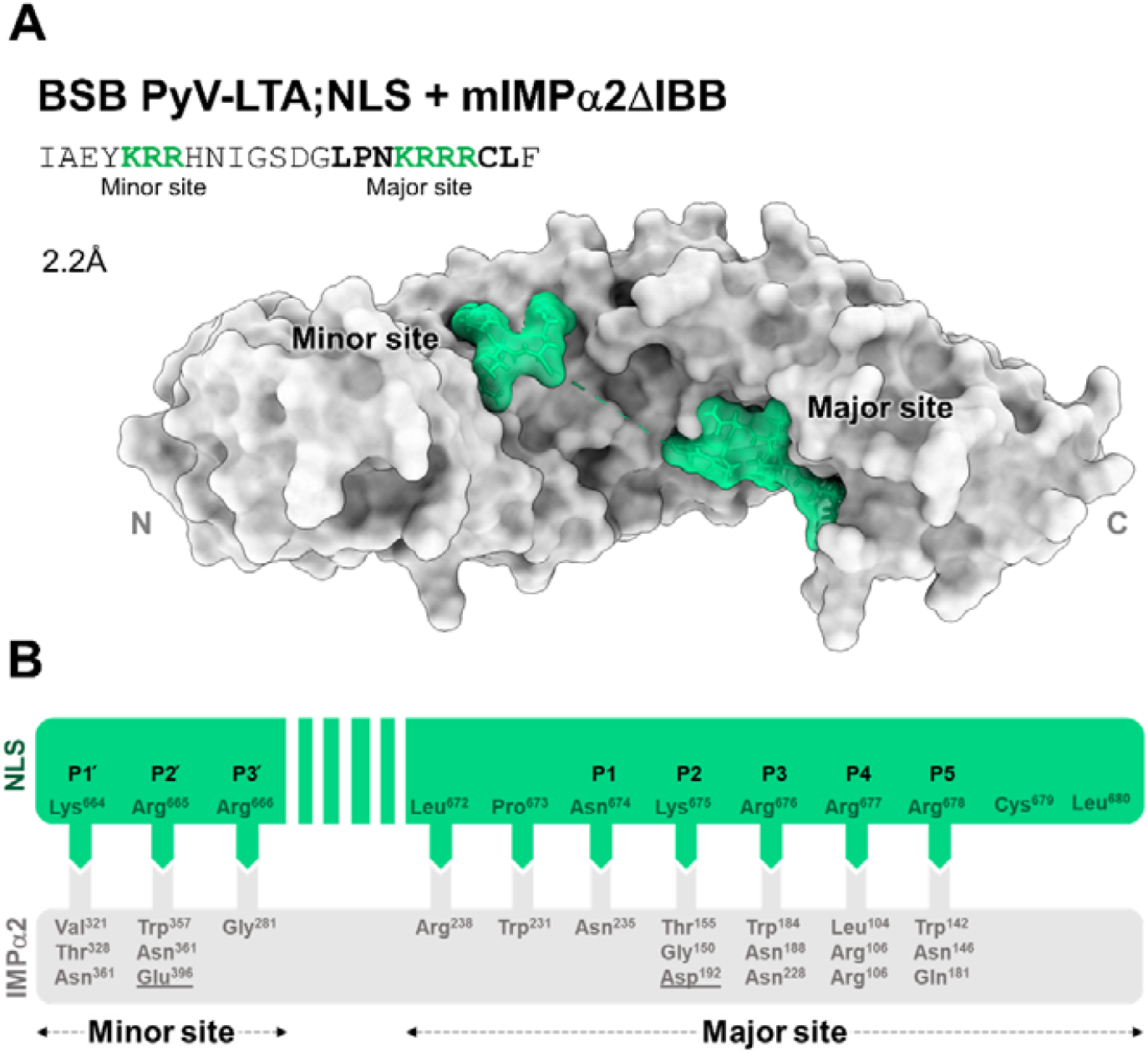
BSB PyV-LTA contains a bipartite NLS at residues 660-683. (A) Crystal structures of the FITC-labelled BSB PyV-LTA;NLS peptide (residues 660-683; green) bound to mIMP ⍰2ΔIBB (gray), with surface representation. (B) Schematic representation of the binding interaction structure, detailing hydrogen bonds and salt bridges (underlined). PDBsum was used for all binding interaction calculations. The structure is deposited with PDB code: 9NIB

### Both basic stretches of amino acids within BSB PyV-LTA bipartite NLS are essential for optimal binding to IMPα2

Since side chains from both basic stretches composing BSB PyV-LTA NLS could be visualized within mIMPα2 binding pockets, we decided to evaluate their contribution to IMP binding. To this end, we assessed the IMP binding ability of FITC-labelled peptides bearing alanine substitution within either the N-terminal (BSB PyV-LTA;mNLSn: 660-IAEYaaaHNIGSDGLPNKRRRCLF-683), the C-terminal (BSB PyV-LTA;mNLSc: 660-IAEYKRRHNIGSDGLPNaaaaCLF-683), or both basic stretches (BSB PyV-LTA;mNLSnc: 660-IAEYaaaHNIGSDGLPNaaaaCLF-683) using EMSA (Figure 5A). Our results demonstrated that both mNLSn and mNLSc could still interact with all IMPs tested, as indicated by co-migration on the native gel, however, a considerable fraction of peptide remained unshifted, suggesting an impaired interaction. On the other hand, incubation of BSB PyV-LTA;mNLSnc peptide with any of the IMPs tested resulted in undetectable co-migration (Figure 5A). These results suggest that both basic stretches of amino acids are important for the interaction of BSB PyV-LTA NLS with IMPs. To better quantify the contribution of each, we also measured the effect of each substitution on binding to mIMPα2ΔIBB by FP (Figure 5B). Our results confirmed that the substitution of both basic stretches to ALA completely impaired mIMPα2ΔIBB binding (Figure 5B). Substitution of the N-terminal decreased binding affinity 10 times, whereas substitution of the C-terminal stretch reduced binding to a higher extent, so that the affinity could not be precisely measured (Figure 5B).

**Figure 5.**
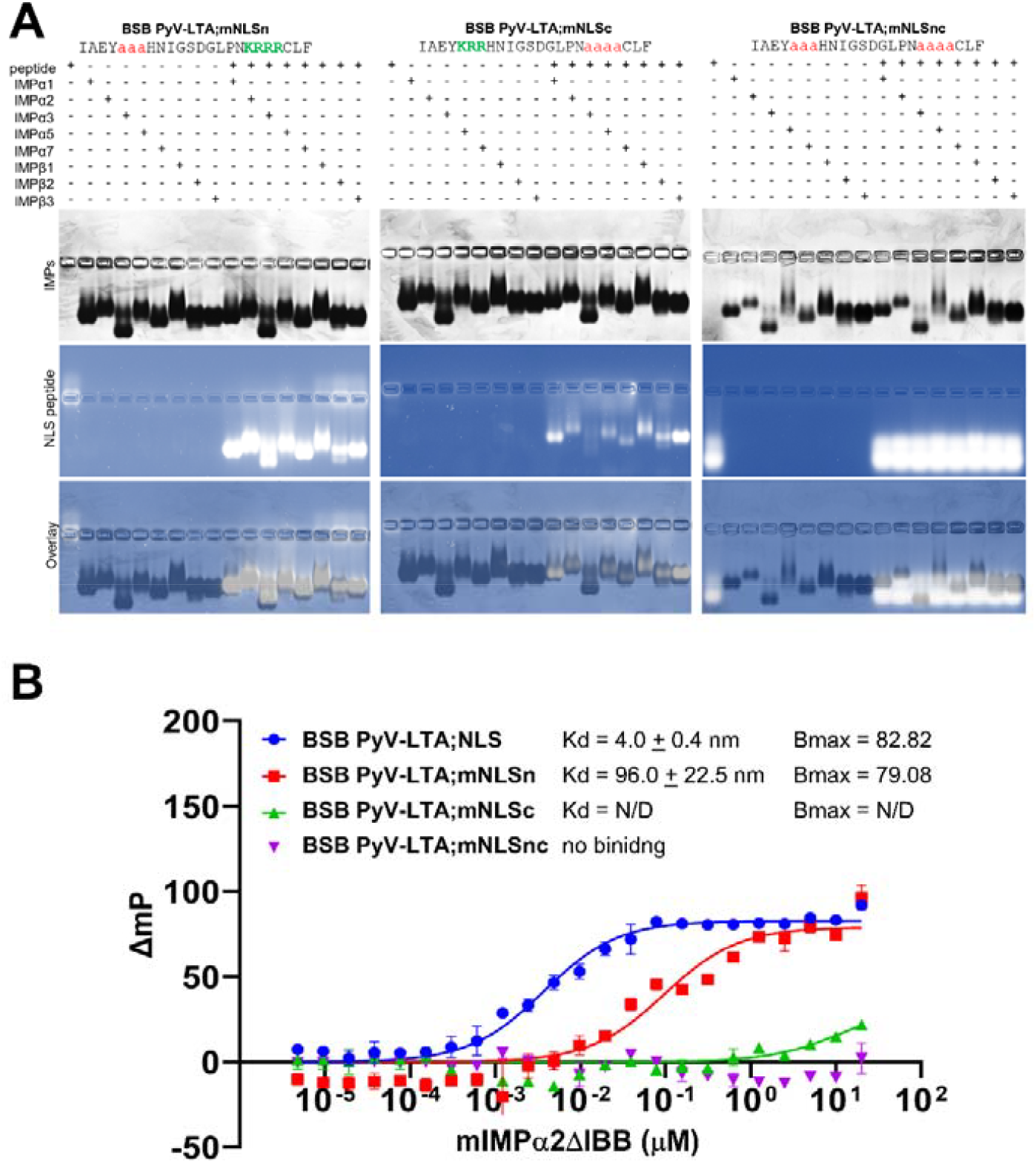
Both stretches of basic amino acids within BSB PyV-LTA bipartite NLS are essential for optimal binding to IMPα. (A) EMSA for FITC labelled BSB PyV-LTA;NLS peptides bearing the indicated alanine substitutions (10 μM) and recombinant IMPs (20 μM). After electrophoresis on 1.5% agarose gel, FITC labelled peptides were visualized with an UV filter (NLS peptides), and proteins by Coomassie staining (IMPs), with an image of both channels shown on bottom (overlay). (B) FP assays were performed by incubating the FITC labelled BSB PyV LTA NLS peptides bearing the indicated alanine substitutions (2 nM) with increasing concentrations of the recombinant mIMPα2ΔIBB, followed by data acquisition and fitting to measure the dissociation constant (Kd) and maximum binding (Bmax) relative to each interaction. Data shown are mean ± standard error of the mean (SEM) relative to three independent experiments.

### Both stretches of basic amino acids within BSB PyV-LTA bipartite NLS are essential for optimal nuclear accumulation

Since BSB PyV-LTA is transported into the nucleus by the IMPα/β1 heterodimer and both stretches of basic amino acids of its cNLS are important for binding to IMPα, we sought to assess their contribution to nuclear transport. To this end, we measured the effect of site-specific substitutions on the levels of nuclear accumulation of BSB PyV LTA-GFP in HEK293A transfected cells by quantitative CLSM (Figure 6A). Substitution of the N-terminal basic residues decreased nuclear accumulation of BSB PyV-LTA-GFP;mNLSn compared to BSB PyV-LTA-GFP (Figure 6B) to a significant extent, with the protein mainly localized to the cytosol in c. 20% of transfected cells (Figure 6C) and a drop of the average Fn/c from 8.8 to 0.8 (Figure 6D). On the other hand, the substitution of the C-terminal stretch of basic amino acids, either alone (BSB PyV-LTA-GFP;mNLSc) or in combination with the N-terminal one (BSB PyV-LTA-GFP;mNLSnc), prevented nuclear accumulation in 100% of transfected cells (Figure 6C), reducing the average Fn/c to 0.3 (Figure 6D). Therefore, both basic stretches of amino acids forming BSB PyV-LTA bipartite NLS are required for nuclear accumulation, and interaction at the IMPα major binding site contributes to a greater extent to IMPα binding and nuclear import.

**Figure 6.**
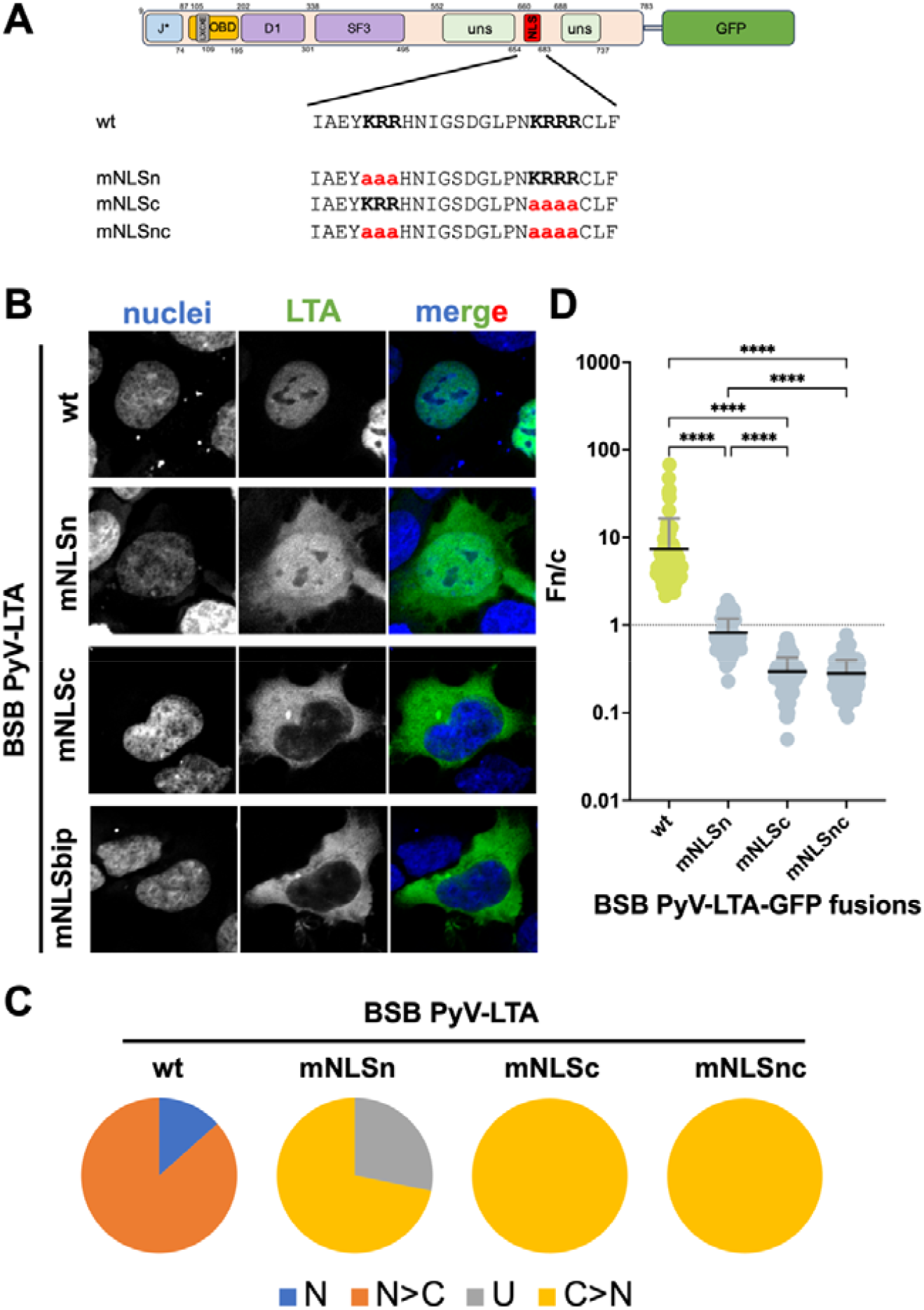
Both stretches of basic amino acids within BSB PyV-LTA bipartite NLS are essential for optimal binding nuclear targeting. (A) Schematic representation of the BSB PyV-LTA-GFP fusion proteins expressed, with indicated the alanine substitutions introduced in each derivative. LXCXE: LXCXE motif; NLS: nuclear localization signal; D1: tag D1 type domain; SF3: SF3 helicase domain; US: unstructured. (B) HEK293A cells were transfected with plasmids encoding the indicated BSB PyV LTA-GFP-fusion derivatives. At 24 h p.t., cells were processed for CLSM and subcellular localization of GFP-fusion proteins was quantitatively analyzed by CLSM. The DRAQ5 (633 nm excitation laser line; nuclei) and the GFP (488 nm excitation laser line; LTA) channels are shown, along with a merged image of the two (merge). (C) Micrographs such as those shown in (B), were quantitively analyzed to calculate the Fn/c relative to individual cells (n > 100). Data shown are the percentage of cells displaying the indicated phenotype. N: Fn/c > 10; N>C: 2< Fn/c < 10; U: 1< Fn/c < 2; C: Fn/c < 1. (D) The mean ± standard deviation of the mean (SD) relative to each GFP-fusion, including the results of the The Welch and Brown-Forsythe one-way ANOVA for significance between the indicated proteins. ****: p < 0.0001.

### cNLS position within LTAs defines the host range of highly divergent PyVs

Despite BSB PyV-LTA possess a cNLS located in a different position compared to LTAs from PyVs infecting mammals, in all cases nuclear transport is mediated by the IMPα/β1 heterodimer (24). This raises the possibility that the cNLSs position in LTA encoded by PyVs is not always conserved, but nuclear localization is preserved. To shed light on this issue, we retrieved from UniProt the coding sequence of the 14 classified PyVs infecting non-mammalian hosts. We also retrieved 5 additional sequences of LTAs from still unclassified novel PyVs recently described from another cartilaginous fish and scorpions (7, 58). We then investigated the conservation of the best characterized LTA domains: the N-terminal J domain along with the HPDKGG hexapeptide; the LXCXE motif; the Tag-D1 type zinc finger domain; the SF3 helicase domain, as well as the presence of putative functional cNLSs, integrating information retrieved from UniProt and ScanProsite with structure-based homology modeling (Supplementary Table V, VI, Supplementary Figure S5-15). Intriguingly, functional domains are heterogeneously conserved among PyVs infecting different hosts (Figure 7AB, Supplementary Figure S16). The least conserved is the LXCXE motif (present in 68.4% of LTAs from PyVs infecting non-mammalian hosts) which is absent in all PyVs infecting fishes, except BSB PyV-LTA. Intriguingly, despite a J domain fold predicted at the N-terminus of all LTAs analyzed, the HPDKGG hexapeptide, essential for Hsp70 interaction, was not identified in LTAs from PyVs infecting ray-finned fishes (Figure 7B, Supplementary Table VI). On the opposite, the OBD, the Tag-D1 type, and the SF3 helicase domains, along with at least a putative cNLS were predicted in all LTAs (Figure 7B). Intriguingly, such cNLSs were detected in extremely variable positions, depending on the infected host (Figure 7C, Supplementary Figure S16). Indeed, 89% of LTAs from PyV infecting birds possess a cNLS within the Tag-D1 type domain, while those infecting scorpions bear a cNLS within the SF3 helicase domain, those infecting ray-finned fishes downstream the SF3 helicase domain, and those infecting cartilaginous fishes within the J domain, respectively (Figure 7C, Supplementary Figure S16, Supplementary Table VI). However, despite only 14% of PyV infecting non-mammalian hosts possessing a cNLS between the LXCXE and the OBD (Figure 7AC, Supplementary Figure S16B), a feature typical of mammalian PyVs (24), a closer inspection of pairwise alignments of LTA sequence from PyVs infecting non-mammalian hosts as compared to SV40-LTA, revealed the presence of an ancestral cNLS corresponding to SV40-LTA NLS in more than in 95% of cases, as shown by the conservation of the TP dipeptide (Figure 7D). This suggests that interhost evolution led to the acquisition of novel cNLS in LTAs from PyVs infecting non-mammalian species, followed by the loss of the cNLS activity from the ancestral NLS. These results support the idea that PyVs have co-speciated with their hosts, acquiring and losing specific domains throughout evolution, based on functional relevance, while IMPα/β1-mediated LTA nuclear localization remained essential for viral replication in all cases.

**Figure 7.**
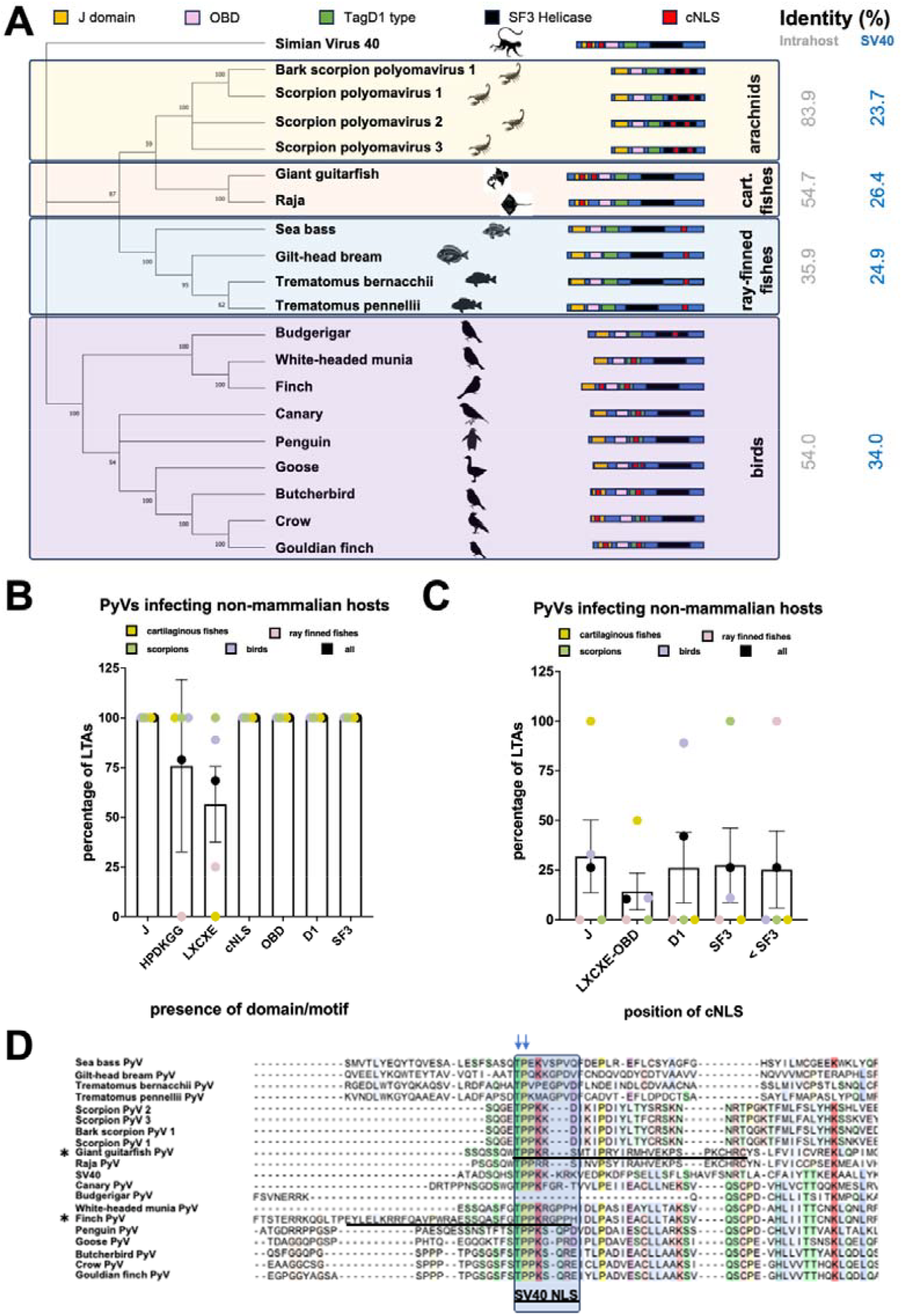
Host-specific cNLSs within LTAs from PyVs infecting non-mammalian hosts. (A) Phylogenetic analysis of LTAs from PyVs infecting non-mammalian hosts rooted on LTA from SV40 as a reference. The position of indicated domains in the protein sequence is shown in the cartoons. The average aminoacidic identity is shown on the right. (B) The percentage of LTAs bearing the specific indicated domain is shown as average ± SD and individual values relative to PyVs infecting non-mammalian hosts. J: J domain; LXCXE: LXCXE motif, D1: Tag-D1 domain; SF3: SF3 helicase domain. (C) The percentage of LTAs bearing a cNLS in the indicated position is shown as average ± SD and individual values relative to PyVs infecting non-mammalian hosts. J: NLS within the J domain; LXCXE-OBD: NLS between the LXCXE motif and the origin binding domain; D1: NLS within the Tag-D1 type domain; SF3: NLS within the SF3 helicase domain; < SF3: NLS downstream of the SF3 helicase domain. (D) The LTA sequences from the indicated PyVs were aligned with clustal Omega. The SV40 LTA cNLS indicated and corresponding sequences are boxed. Vertical arrows indicate the conserved TP dipeptide. The cNLSs from Giant guitarfish PyV and Finch PyV LTA, identified by cNLS mapper as functional, are underlined.

## DISCUSSION

The high sequence diversity of PyVs LTA, combined with their necessity to localize in the nucleus of mediate viral replication, provides an exceptional opportunity to study the evolution of their host-cell interactions. Our analysis indicates that while an OBD, a helicase domain, a Tag-D1 type motif and a cNLS are always present, other functional domains and motifs correlate with PyVs host species (Figure 7AB). This suggests that all LTAs originate from a common ancestral precursor with nuclear targeting and DNA helicase properties and subsequently evolved to perform additional species-specific functions (58). The LXCXE motif is absent in LTAs from PyVs infecting either ray-finned or cartilaginous fishes – apart from BSB PyV, while the HPDKGG hexapeptide motif is absent exclusively in LTAs from PyVs infecting ray-finned fishes (Figure 7B, Supplementary Table VI). This suggests that the ability to interact with host chaperone proteins as mediated by a functional J domain (12) emerged before the ability to inactivate Rb (13). Our data also strongly implies functional convergence of all LTAs on the IMPα/β1 pathway to reach the cell nucleus. This is supported by the bioinformatics identification of putative cNLS in LTAs from all known PyVs (Table I, Supplementary Table V), and by a detailed characterization of nuclear import of BSB PyV-LTA, which is less than 30% identical to SV40-LTA. We provide compelling evidence that BSB PyV-LTA can localize in the cell nucleus independently from any other viral protein and that the process can be hindered by the well-known IMPα/β1 inhibitor Bimax2 (Figure 3), analogously to all other LTAs tested so far (24). Such observation is intriguing since several alternative nuclear import pathways appear highly conserved in eukaryotic cells (59). The reason for such preference for IMPα/β1 dependent nuclear import might reflect the fact that it has an increased dynamic range for control of import rates and a more flexible control of cargo gradients under different cellular conditions, resulting in increased robustness against environmental influences as compared to other nuclear import pathways (60). Intriguingly, BSB PyV-LTA NLS appears to be bipartite, being able to simultaneously bind to both mIMPα2ΔIBB minor and major sites (Figure 4), similarly to several other LTAs, such as those encoded by Saint Louis and KI Polyomaviruses, but in stark contrast to SV40, JC, and BK PyVs (24). Since cargo bearing bipartite NLSs can accumulate in the nucleus to a higher concentration than those bearing a monopartite NLS (60), this could reflect the importance of LTA nuclear localization for BSB PyVs replication. As observed for several other bipartite NLSs (24, 61–66), both basic stretches of amino acids are required for IMPα binding (Figure 5) and nuclear accumulation (Figure 6), despite the downstream sequence, directly interacting with IMPα major binding site, contributes to a higher extent to both processes. Additionally, the observation that cNLS position within LTAs is dependent on the infected host (Figure 7C) might help the classification of novel-identified PyVs. Indeed, the still unclassified Giant Guitarfish PyV encodes an LTA which shares 54.7% identity with etapolyomavirus Raja PyV LTA (Figure 7A), and they both possess a putative cNLS in the J domain (Figure 7AC, Supplementary Figures S10, S11 and S16H) so that they could both be classified in the same genus. On the same lines, all LTAs from scorpions share 83.9% identity and can be distinguished by all others by the presence of a putative cNLS within the Tag-D1 motif, suggesting they should be classified in a separate genus. Finally, the presence of ancestral cNLSs located upstream of the OBD in all LTAs analyzed (Figure 7D), implies that PyV LTAs share a common ancestor endowed with a cNLS upstream of the OBD.

**Table I.**
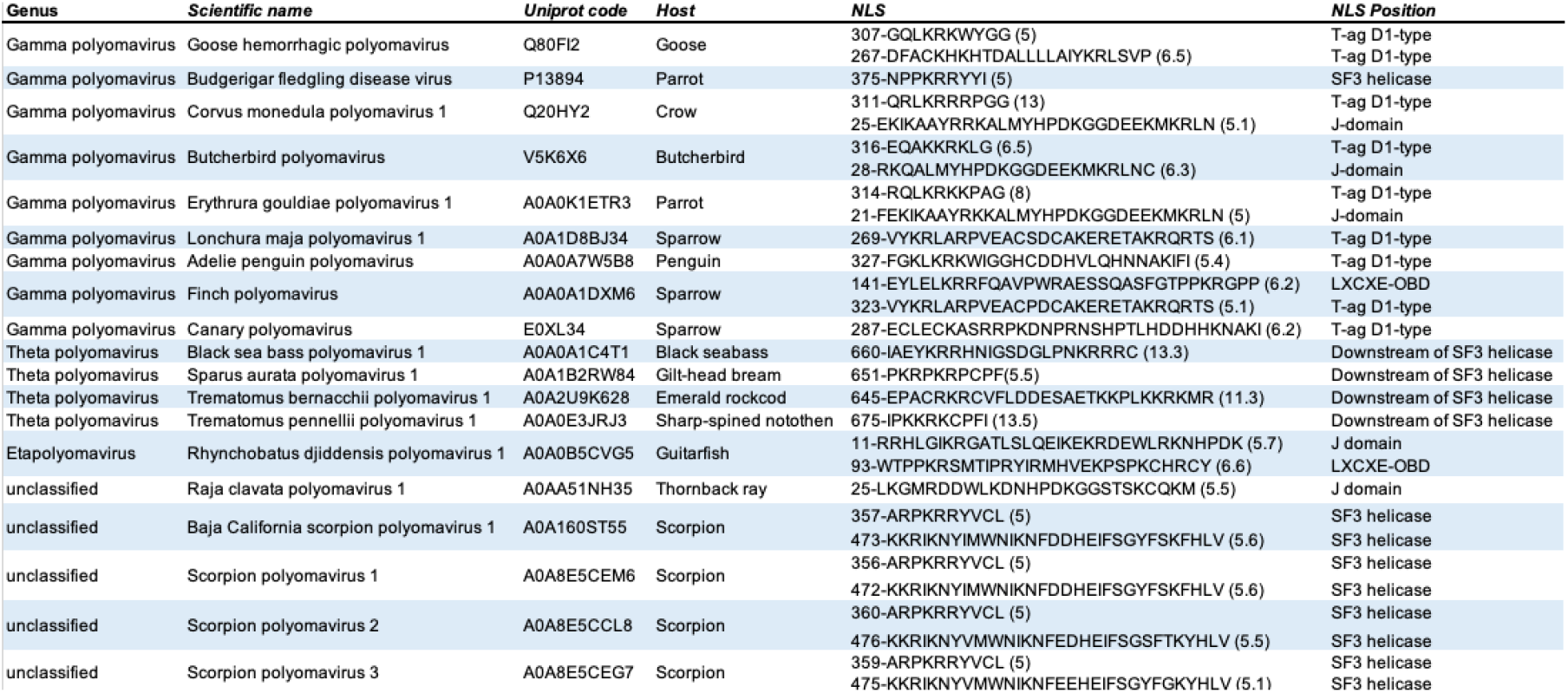
Putative cNLSs in LTA from PyVs infecting non-mammalian hosts. Details of each PyV infecting non-mammalian hosts analyzed are provided, such as genus and virus scientific name and infected host. The UniProt code of each LTA is also shown, along with NLS position, sequence, cNLS score and position.

## Author Contributions

Conceptualization: G.A.; methodology: O.T., J.R.; validation: G.A., M.H.; formal analysis: G.A., M.H., S.N., S.P.; Investigation: G.A., M.H., S.N., S.P.; data curation: G.A., M.H., G.A.; writing— original draft preparation: G.A., M.H.; writing—review and editing: G.A., M.H., S.N., S.P., J.F., O.T., J.R.; visualization: G.A., M.H.; supervision: G.A.; project administration: G.A., J.F.; funding acquisition: G.A. All authors have read and agreed to the published version of the manuscript.

## Acknowledgements

This research was undertaken in part using the MX2 beamline at the Australian Synchrotron, part of ANSTO, and made use of the Australian Cancer Research Foundation (ACRF) detector. We thank Christopher Buck (Bethesda) for helpful discussions and preliminary revision of the manuscript.

## Supplementary Material

The following supplementary Material is available online: Supplementary Figure S1 analytical data for production of BSB PyV-LTA;NLS peptide. Supplementary Figure S2 analytical data for production of BSB PyV-LTA;mNLSn peptide. Supplementary Figure S3 analytical data for production of BSB PyV-LTA;mNLSc peptide. Supplementary Figure S4 analytical data for production of BSB PyV-LTA;mNLSnc. Supplementary Figures S5-S15 AlphaFold models for LTA from PyVs infecting fishes and scorpions. Supplementary Figure S16. Presence and position of functional domains and motifs in LTAs from PyVs infecting non-mammalian hosts. Supplementary Table I. List of plasmids used in this study. Supplementary Table II. List of peptides used in this study. Supplementary Table III. Crystallization information and data processing statistics. Supplementary Table IV. Calculated interactions of BSB PyV-LTA NLS and IMP ⍰2ΔIBB structure. Supplementary Table V. Details of LTAs from PyVs infecting non-mammalian hosts. Supplementary Table VI. Functional domains and motifs identifiied in LTAs from PyVs infecting non-mammalian hosts.

