## Supplementary Figure for "Importin α/β1 dependent nuclear import of Black Sea Bass Polyomavirus Large Tumor Antigen is mediated by a classical NLS located downstream of the SF3 helicase domain"

**Positioning of Nuclear Localization Signals as an Evolutionary Marker in Polyomavirus Large Tumor Antigens**

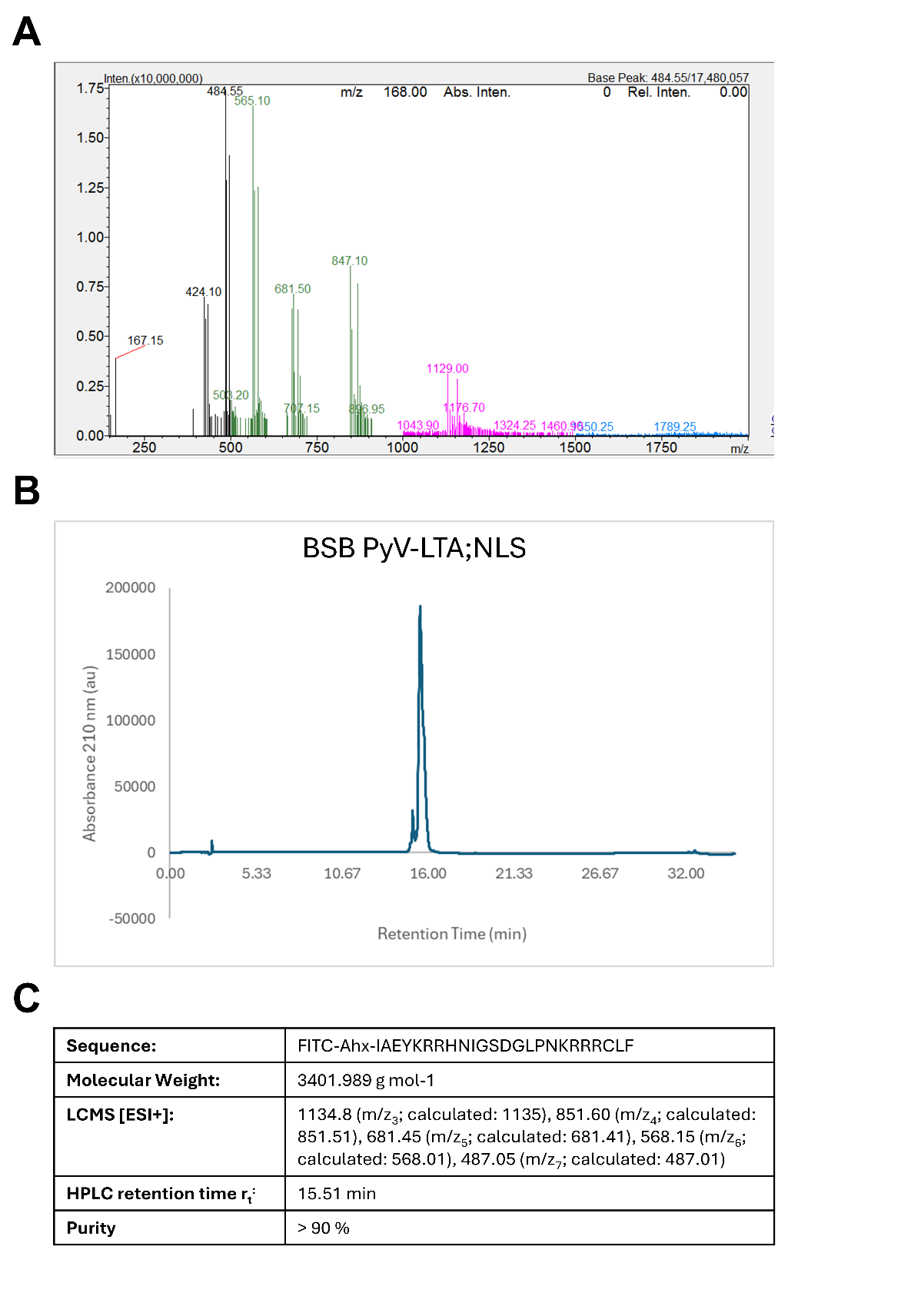

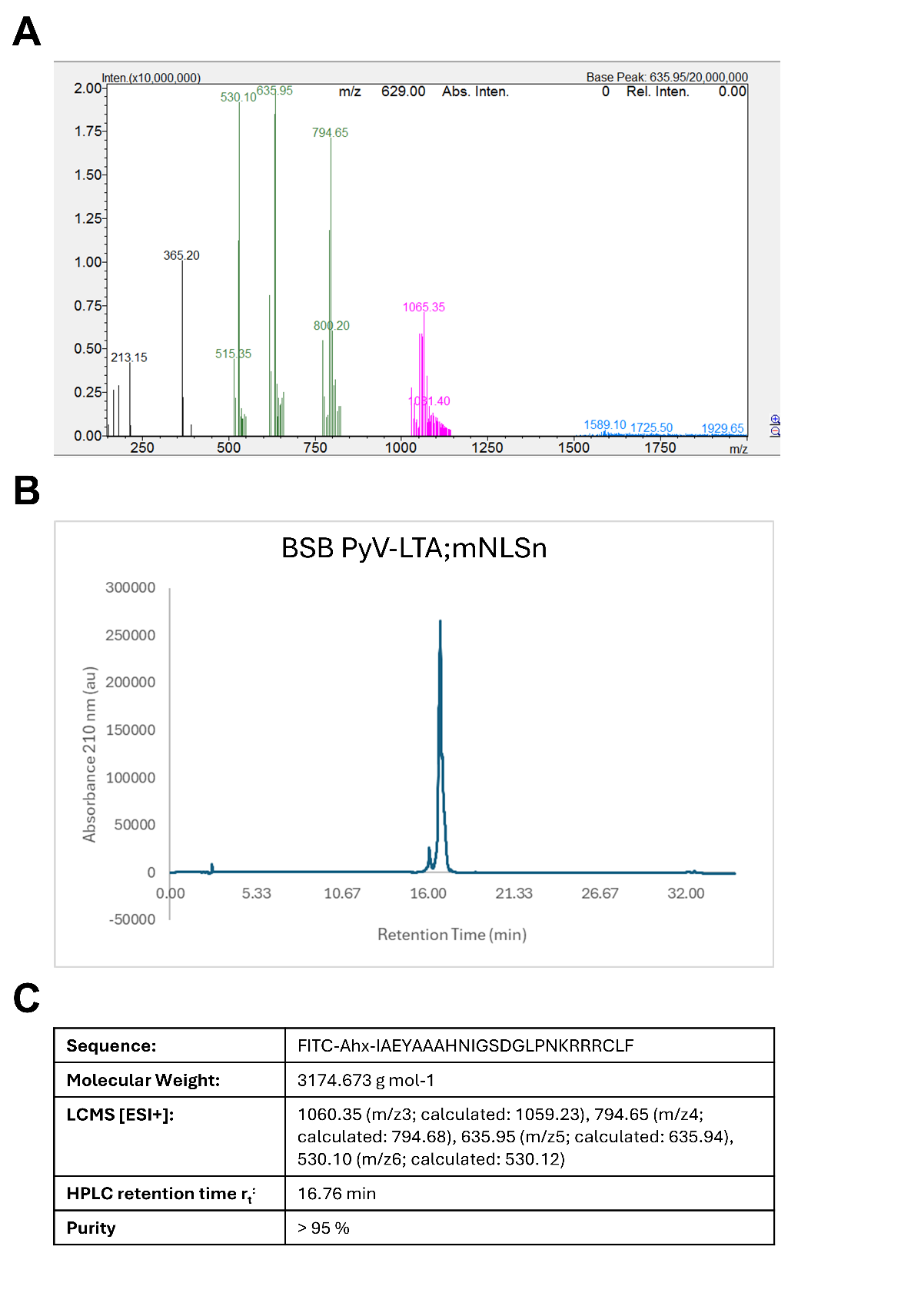

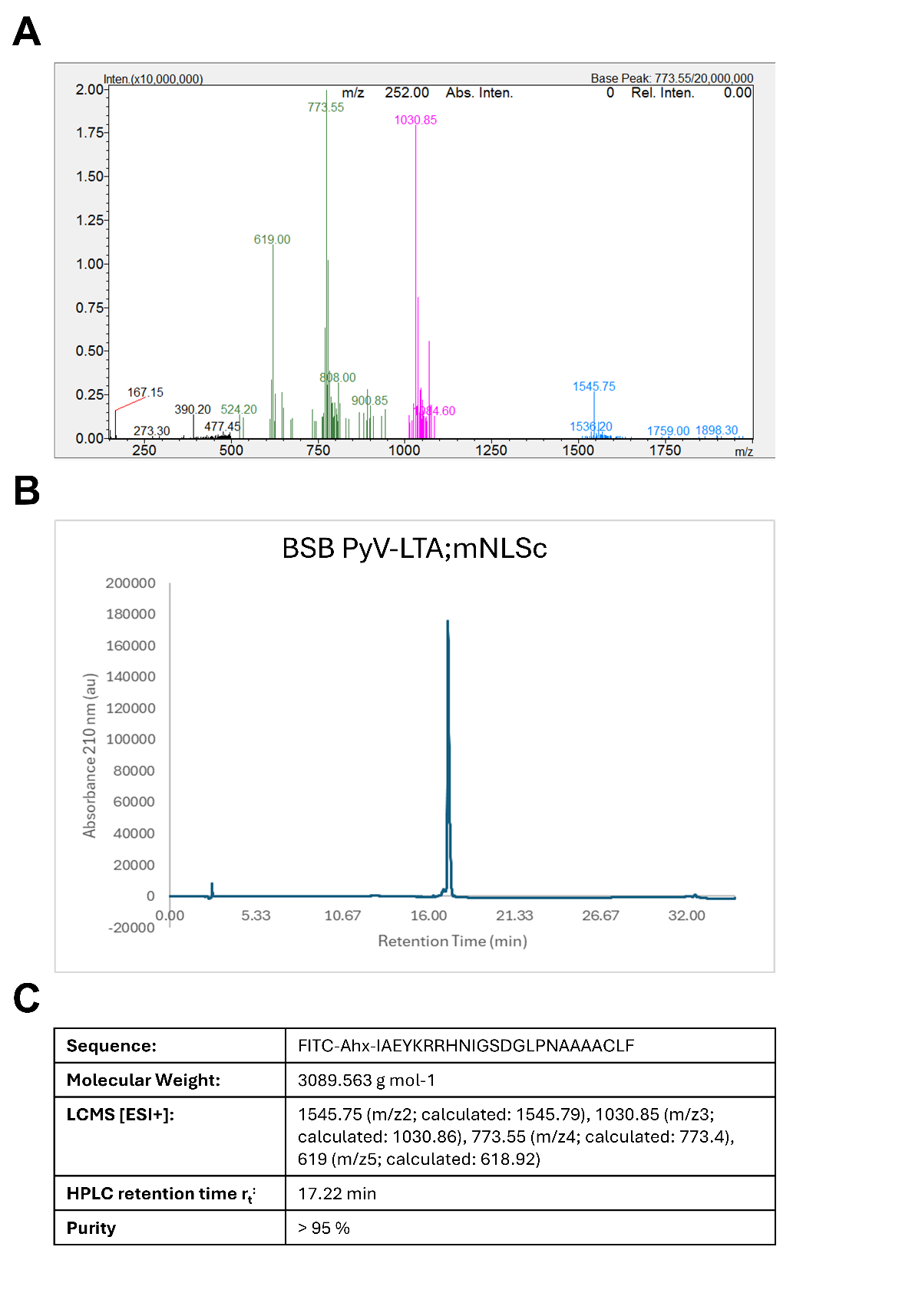

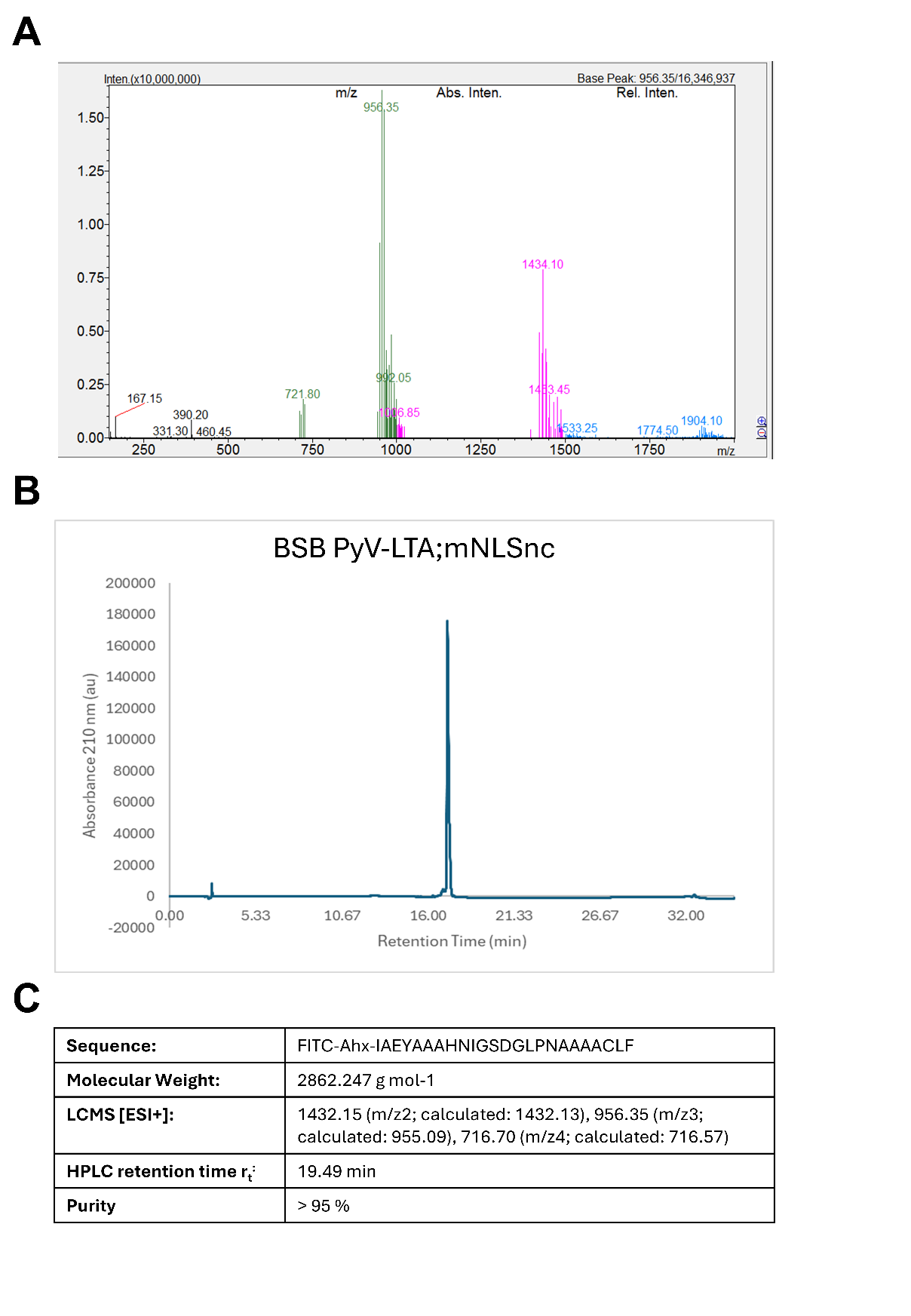

**Supplementary Figure S1. Analytical data from production of peptide BSB PyV-LTA;NLS.** (A) LCMS trace (B) HPLC chromatogram (C) Analytical data summary.

**Supplementary Figure S2. Analytical data from production of peptide BSB PyV-LTA;NLSn.** (A) LCMS trace (B) HPLC chromatogram (C) Analytical data summary.

**Supplementary Figure S3. Analytical data from production of peptide BSB PyV-LTA;NLSc.** (A) LCMS trace (B) HPLC chromatogram (C) Analytical data summary.

**Supplementary Figure S4. Analytical data from production of peptide BSB PyV-LTA;NLSnc.** (A) LCMS trace (B) HPLC chromatogram (C) Analytical data summary.

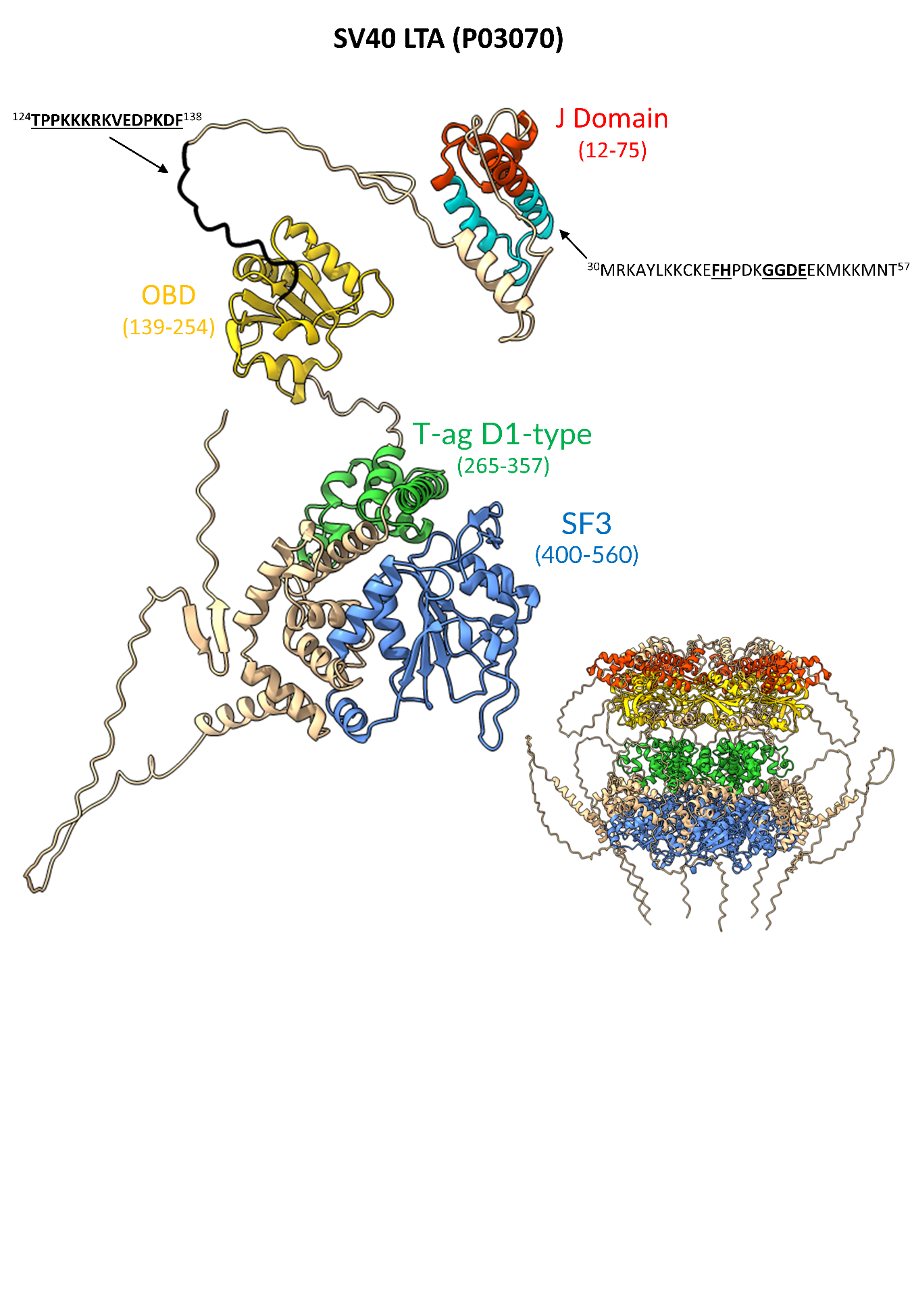
**Supplementary Figure S5. Structural model of SV40 LTA and functional domains.** Monomeric and hexameric forms of SV40 LTA protein (UniProt code: P03070) predicted using AlphaFold3. The specific domains and motifs are colored according to the following scheme: J domain, red; origin-binding domain (OBD), yellow; T-ag D1 type, green; helicase domain (SF3), blue. Residues encompassing these domains are shown alongside the domain label in parentheses. The identified cNLSs have been indicated with arrows, and colored according to the scheme: residues 30-57 (cNLS within J domain), cyan; residues 124-138 (cNLS upstream of the OBD), black. cNLS sequences are shown with residues in bold and underlined representing flexible regions of the structure whilst those not bold or underlined are identified to be part of the predicted secondary structures.

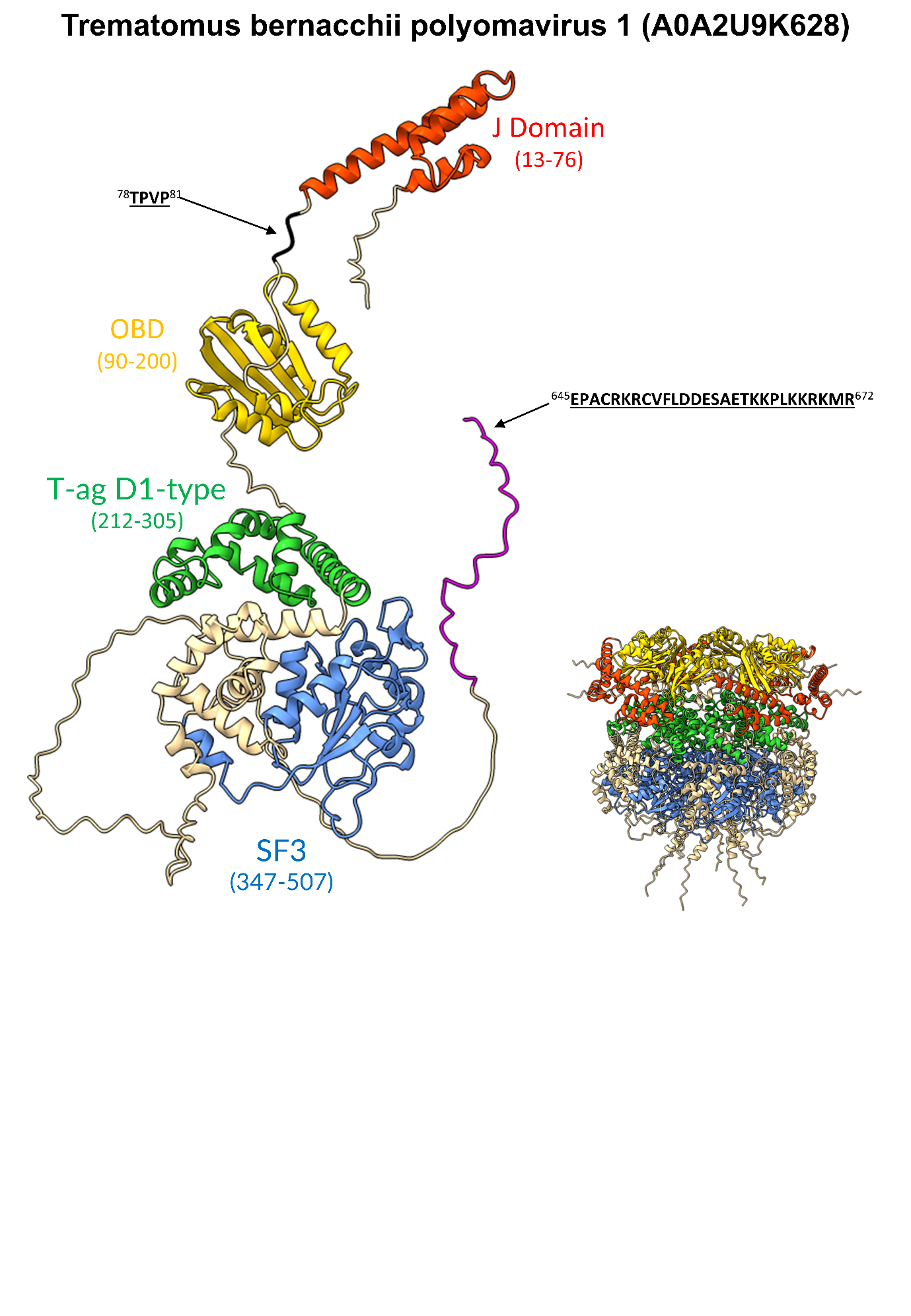

**Supplementary Figure S6. Structural model of Trematomus bernacchi polyomavirus 1 LTA functional domains.** Monomeric and hexameric forms of Trematomus bernacchi PyV 1 LTA protein (UniProt code: A0A2U9K628) predicted using AlphaFold3. The specific domains and motifs are colored according to the following scheme: J domain, red; origin-binding domain (OBD), yellow; T-ag D1 type, green; helicase domain (SF3), blue. Residues encompassing these domains are shown alongside the domain label in parentheses. The identified cNLSs have been indicated with arrows, and colored according to the scheme: residues 654-672 (cNLS downstream of the SF3 domain), magenta; residues 78-81 (ancestral cNLS upstream of the OBD), black. cNLS sequences are shown with residues in bold and underlined representing flexible regions of the structure whilst those not bold or underlined are identified to be part of the predicted secondary structures.

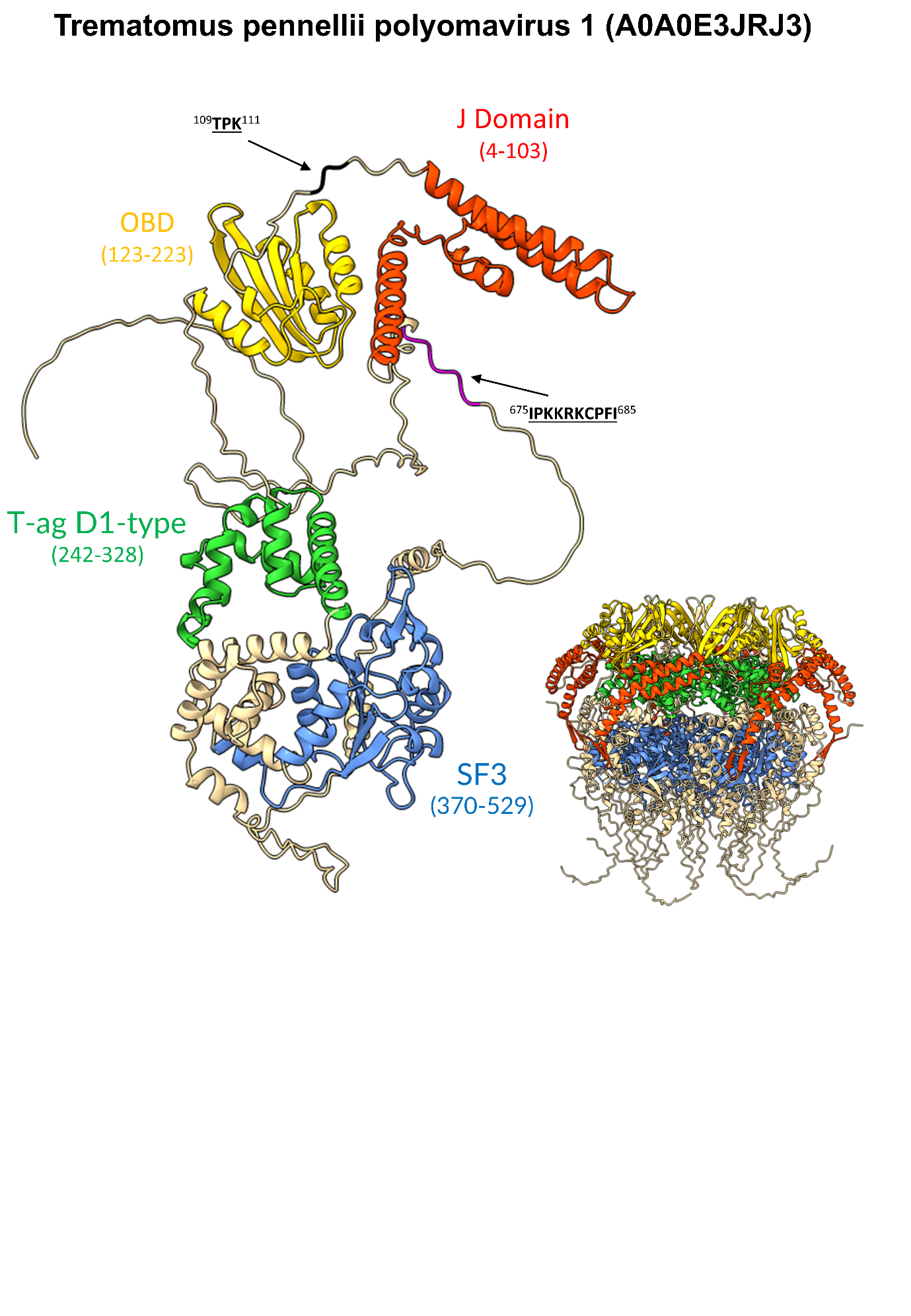

**Supplementary Figure S7. Structural model of Trematomus pennellii polyomavirus 1 LTA functional domains.** Monomeric and hexameric forms of Trematomus pennellii PyV 1 LTA protein (UniProt code: A0A0E3JRJ3) predicted using AlphaFold3. The specific domains and motifs are colored according to the following scheme: J domain, red; origin-binding domain (OBD), yellow; T-ag D1 type, green; helicase domain (SF3), blue. Residues encompassing these domains are shown alongside the domain label in parentheses. The identified cNLSs have been indicated with arrows, and colored according to the scheme: residues 675-685 (cNLS downstream of the SF3 domain), magenta; residues 109-111 (ancestral cNLS upstream of the OBD), black. cNLS sequences are shown with residues in bold and underlined representing flexible regions of the structure whilst those not bold or underlined are identified to be part of the predicted secondary structures.

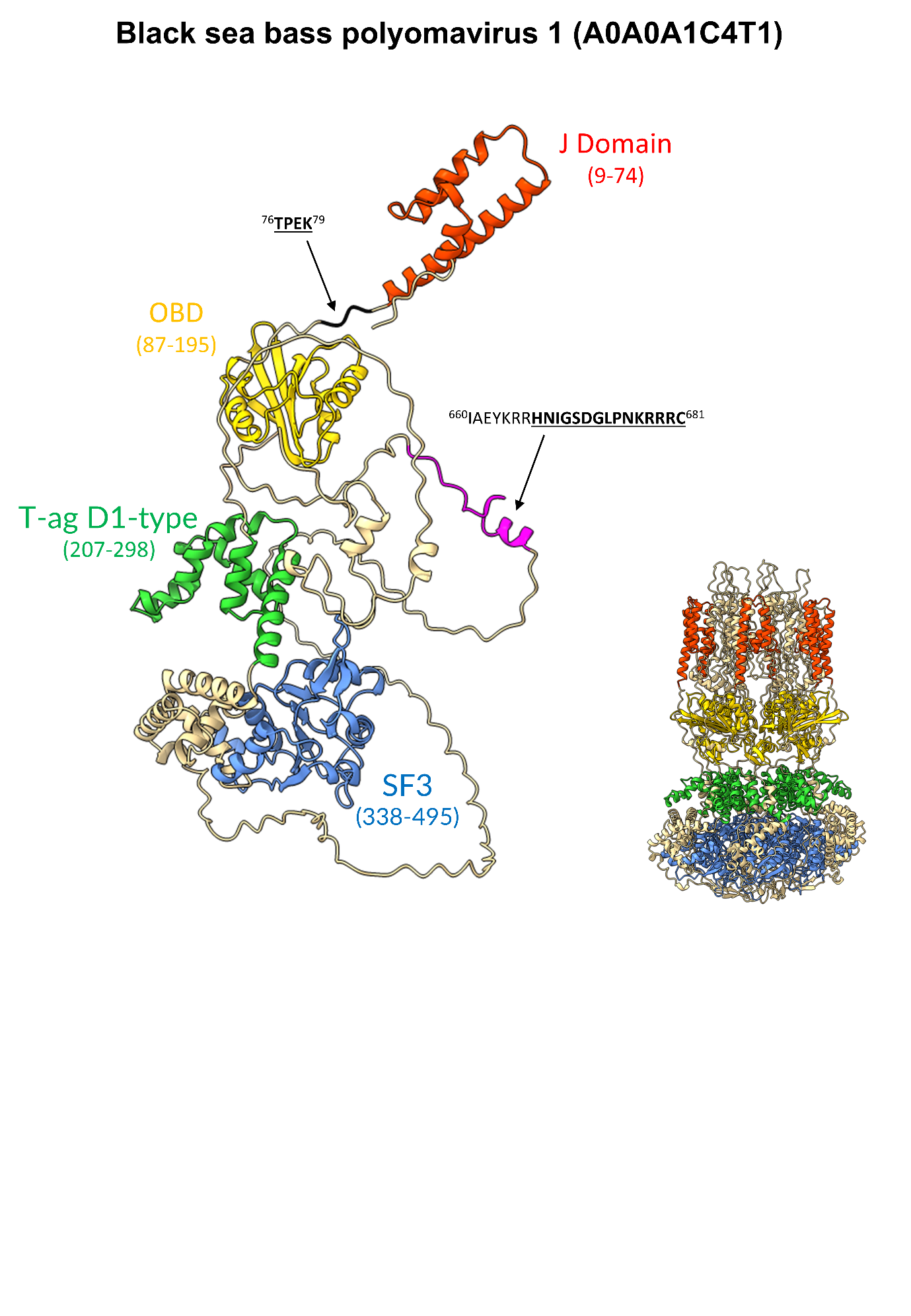
**Supplementary Figure S8. Structural model of Black Sea Bass polyomavirus 1 LTA functional domains.** Monomeric and hexameric forms of Black sea bass PyV1 LTA protein (UniProt code: A0A0A1C4T1) predicted using AlphaFold3. The specific domains and motifs are colored according to the following scheme: J domain, red; origin-binding domain (OBD), yellow; T-ag D1 type, green; helicase domain (SF3), blue. Residues encompassing these domains are shown alongside the domain label in parentheses. The identified cNLSs have been indicated with arrows, and colored according to the scheme: residues 660-681 (cNLS downstream of the SF3 domain), magenta; residues 76-79 (ancestral cNLS upstream of the OBD), black. cNLS sequences are shown with residues in bold and underlined representing flexible regions of the structure whilst those not bold or underlined are identified to be part of the predicted secondary structures.

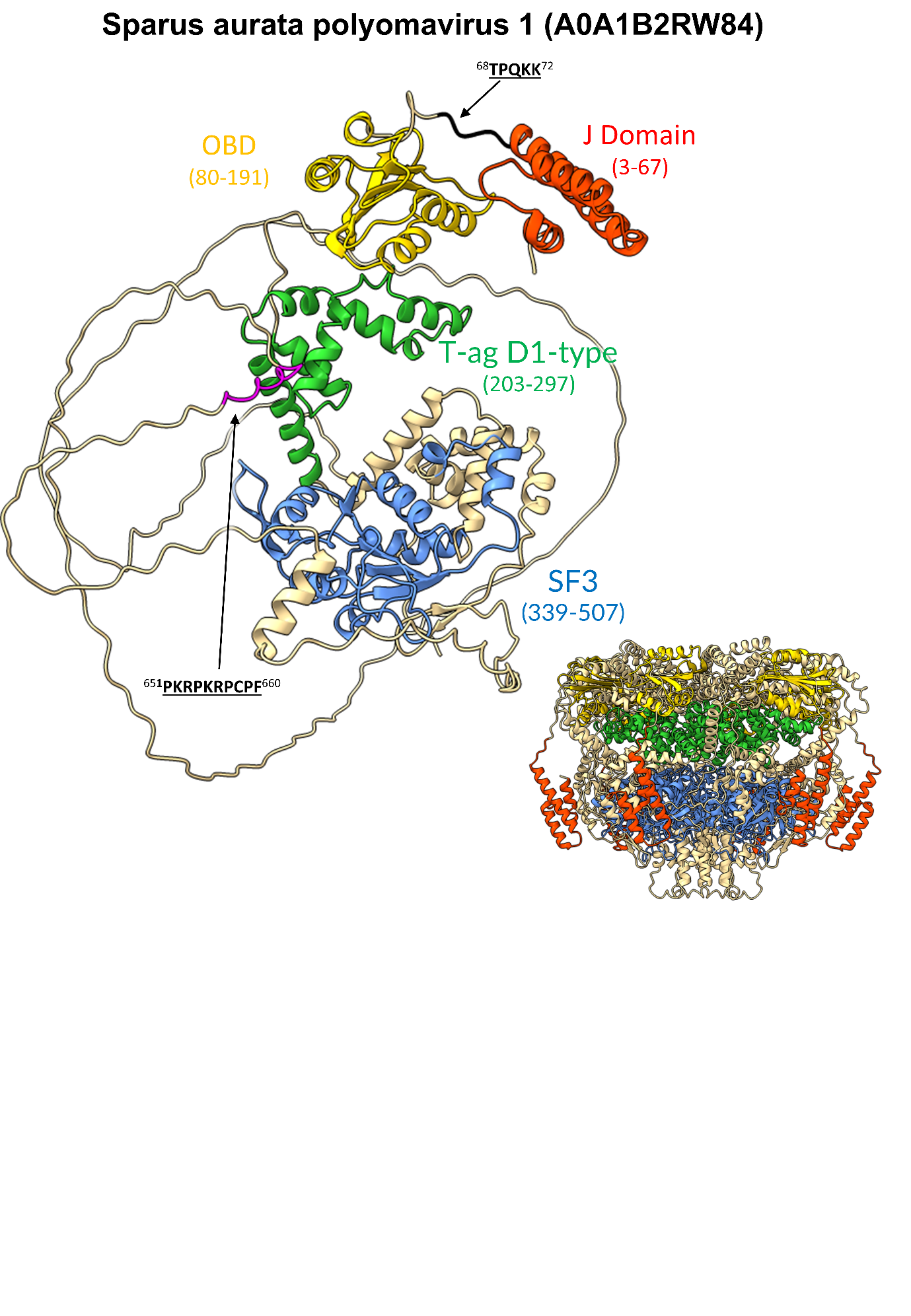
**Supplementary Figure S9. Structural model of Sparus aurata polyomavirus 1 LTA functional domains.** Monomeric and hexameric forms of Sparus aurata PyV 1 LTA protein (UniProt code: A0A1B2RW84) predicted using AlphaFold3. The specific domains and motifs are colored according to the following scheme: J domain, red; origin-binding domain (OBD), yellow; T-ag D1 type, green; helicase domain (SF3), blue. Residues encompassing these domains are shown alongside the domain label in parentheses. The identified cNLSs have been indicated with arrows, and colored according to the scheme: residues 651-660 (cNLS downstream of the SF3 domain), magenta; residues 68-72 (ancestral cNLS upstream of the OBD), black. cNLS sequences are shown with residues in bold and underlined representing flexible regions of the structure whilst those not bold or underlined are identified to be part of the predicted secondary structures.

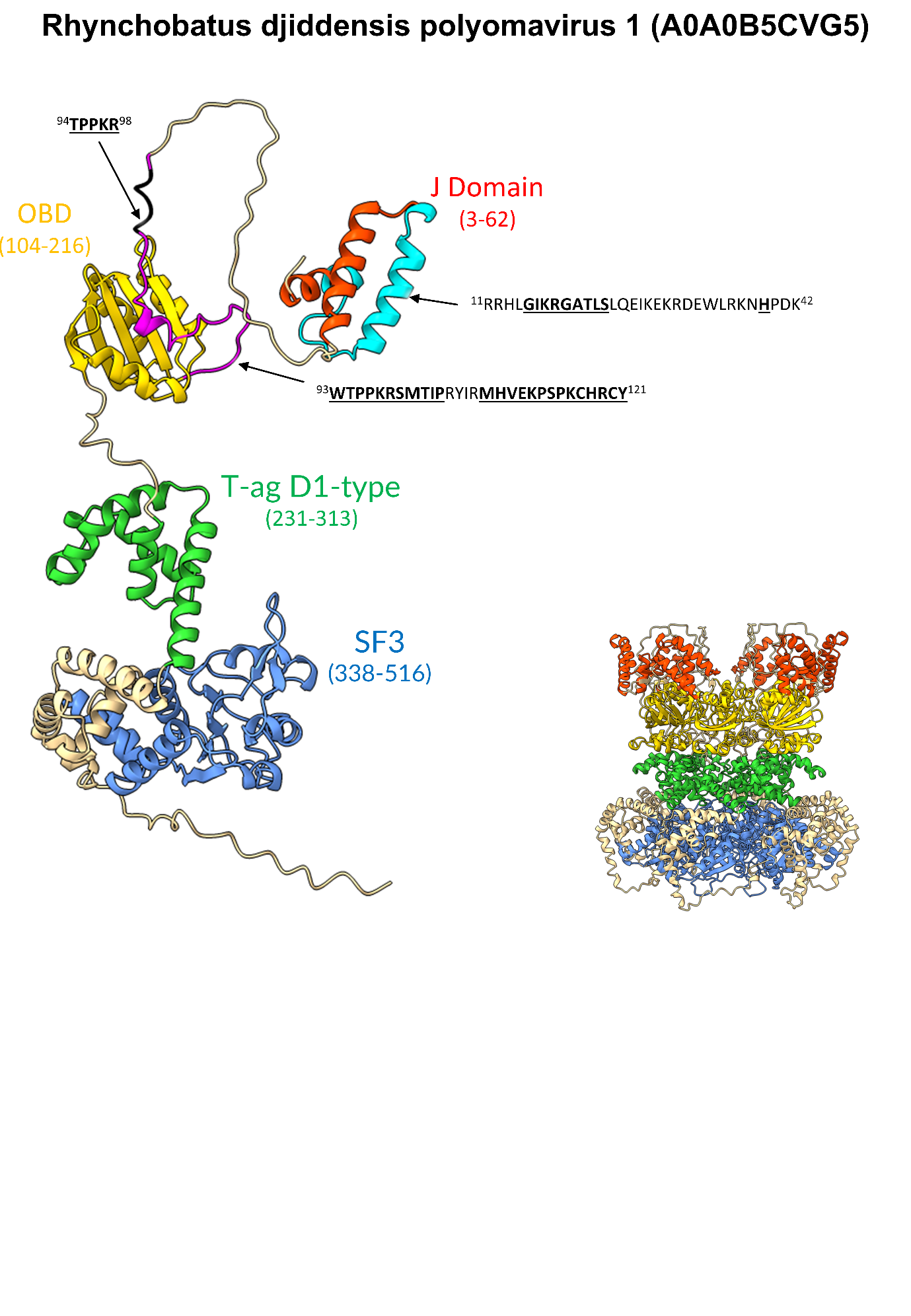

**Supplementary Figure S10. Structural model of Rhynchobatus djiddensis polyomavirus 1 LTA functional domains.** Monomeric and hexameric forms of Rhynchobatus djiddensis PyV 1 LTA protein (UniProt code: A0A0B5CVG5) predicted using AlphaFold3. The specific domains and motifs are colored according to the following scheme: J domain, red; origin-binding domain (OBD), yellow; T-ag D1 type, green; helicase domain (SF3), blue. Residues encompassing these domains are shown alongside the domain label in parentheses. The identified cNLSs have been indicated with arrows, and colored according to the scheme: residues 11-42 (cNLS within the J domain), cyan; residues 93-121 (NLS upstream of OBD) magenta; residues 94-98 (ancestral cNLS upstream of the OBD, overlaps magenta NLS), black. cNLS sequences are shown with residues in bold and underlined representing flexible regions of the structure whilst those not bold or underlined are identified to be part of the predicted secondary structures.

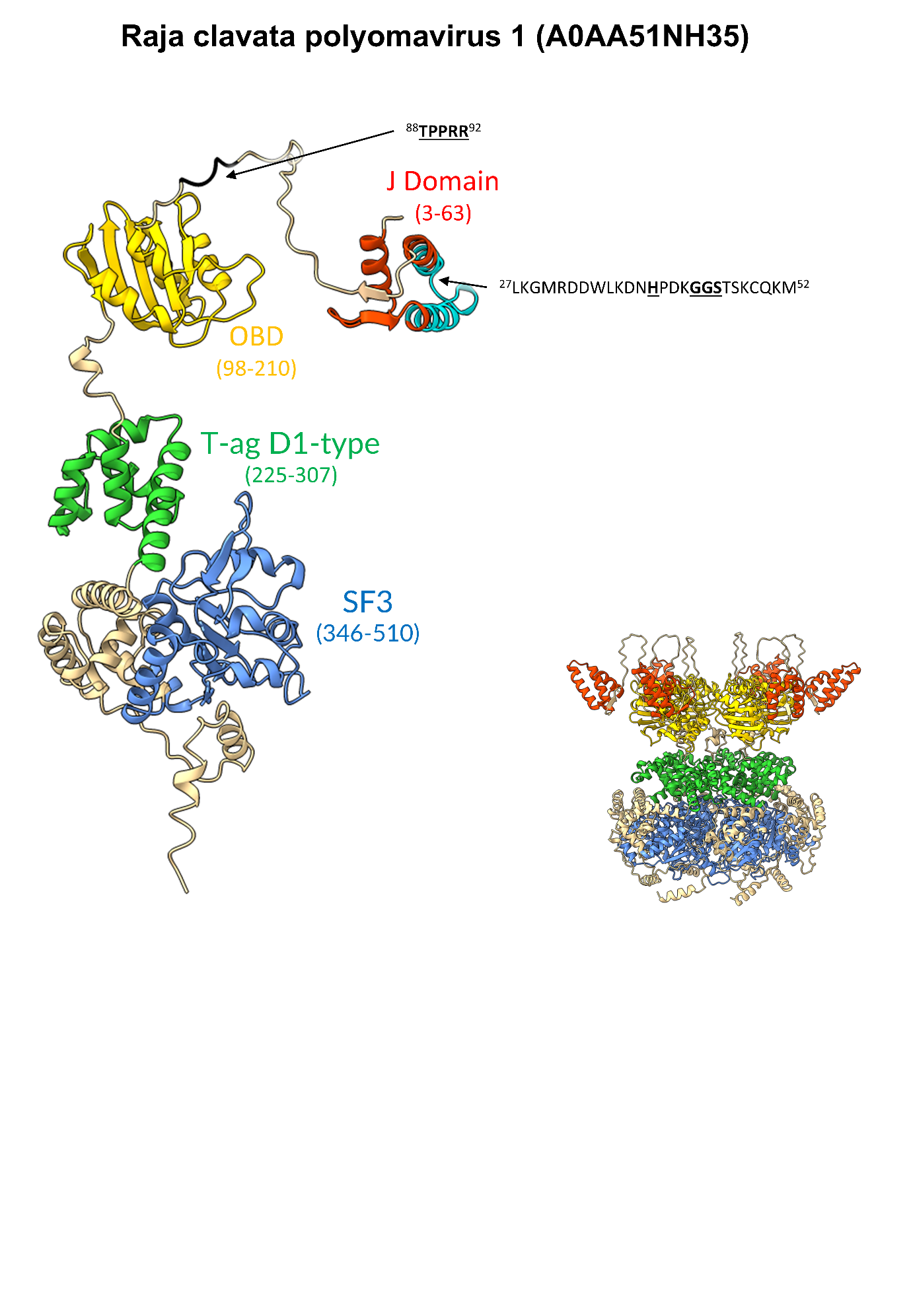

**Supplementary Figure S11. Structural model of Raja clavata polyomavirus 1 LTA functional domains.** Monomeric and hexameric forms of Raja clavate PyV 1 LTA protein (UniProt code: A0AA51NH35) predicted using AlphaFold3. The specific domains and motifs are colored according to the following scheme: J domain, red; origin-binding domain (OBD), yellow; T-ag D1 type, green; helicase domain (SF3), blue. Residues encompassing these domains are shown alongside the domain label in parentheses. The identified cNLSs have been indicated with arrows, and colored according to the scheme: residues 27-52 (cNLS within the J domain), cyan; residues 88-92 (ancestral cNLS upstream of the OBD), black. cNLS sequences are shown with residues in bold and underlined representing flexible regions of the structure whilst those not bold or underlined are identified to be part of the predicted secondary structures.

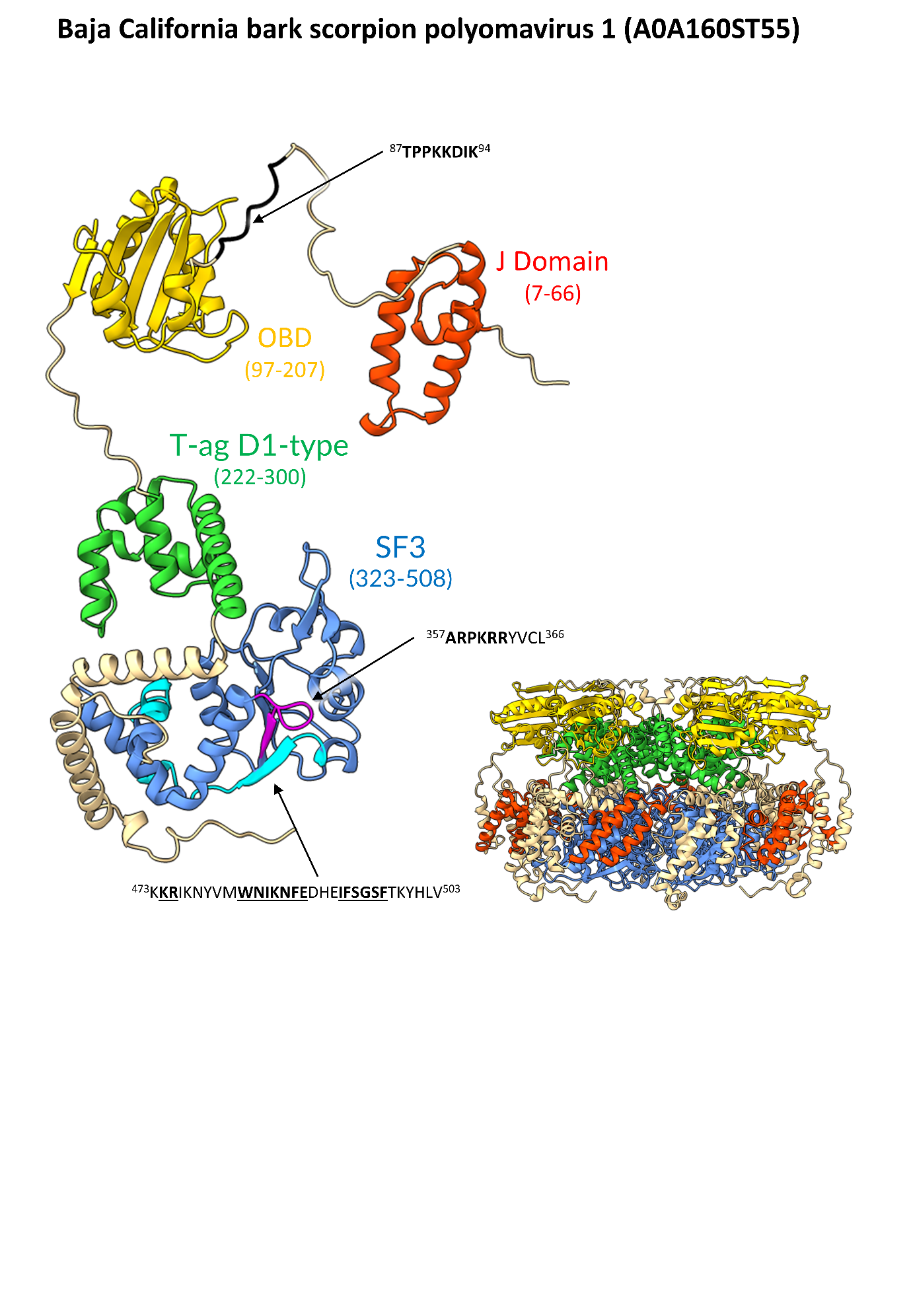

**Supplementary Figure S12. Structural model of Baja California bark scorpion polyomavirus 1 LTA functional domains.** Monomeric and hexameric forms of Baja California black scorpion PyV 1 LTA protein (UniProt code: A0A160ST55) predicted using AlphaFold3. The specific domains and motifs are colored according to the following scheme: J domain, red; origin-binding domain (OBD), yellow; T-ag D1 type, green; helicase domain (SF3), blue. Residues encompassing these domains are shown alongside the domain label in parentheses. The identified cNLSs have been indicated with arrows, and colored according to the scheme: residues 357-366 (cNLS within the SF3 domain), magenta; residues 473-503 (cNLS within the SF3 domain), cyan; residues 87-94 (ancestral cNLS upstream of the OBD), black. cNLS sequences are shown with residues in bold and underlined representing flexible regions of the structure whilst those not bold or underlined are identified to be part of the predicted secondary structures.

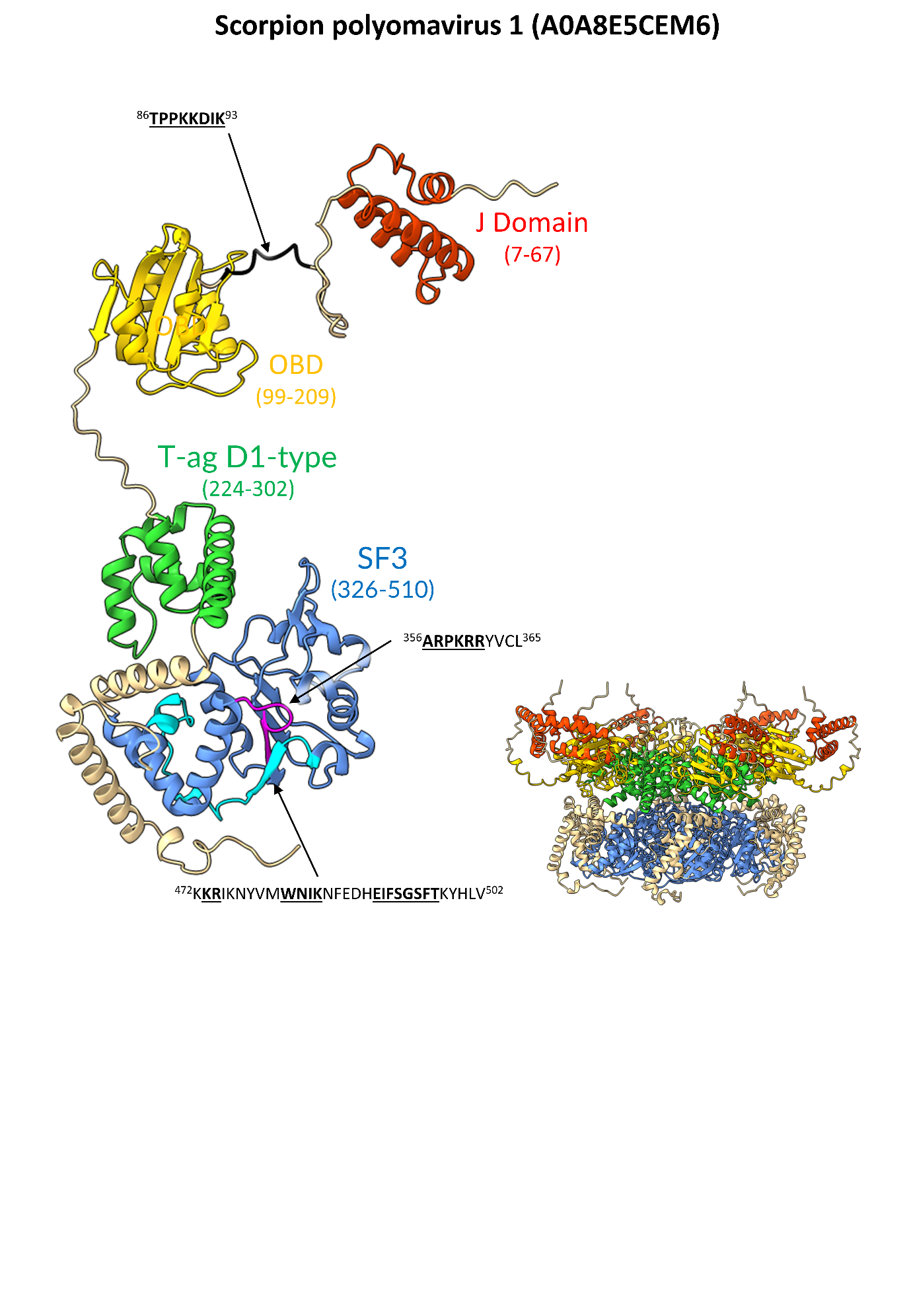

**Supplementary Figure S13. Structural model of Scorpion polyomavirus 1 LTA functional domains.** Monomeric and hexameric forms of scorpion PyV 1 LTA protein (UniProt code: A0A8E5CEM6) predicted using AlphaFold3. The specific domains and motifs are coloured according to the following scheme: J domain, red; origin-binding domain (OBD), yellow; T-ag D1 type, green; helicase domain (SF3), blue. Residues encompassing these domains are shown alongside the domain label in parentheses. The identified cNLSs have been indicated with arrows, and colored according to the scheme: residues 356-365 (cNLS within the SF3 domain), magenta; residues 472-502 (cNLS within the SF3 domain), cyan; residues 86-93 (ancestral cNLS upstream of the OBD), black. cNLS sequences are shown with residues in bold and underlined representing flexible regions of the structure whilst those not bold or underlined are identified to be part of the predicted secondary structures.

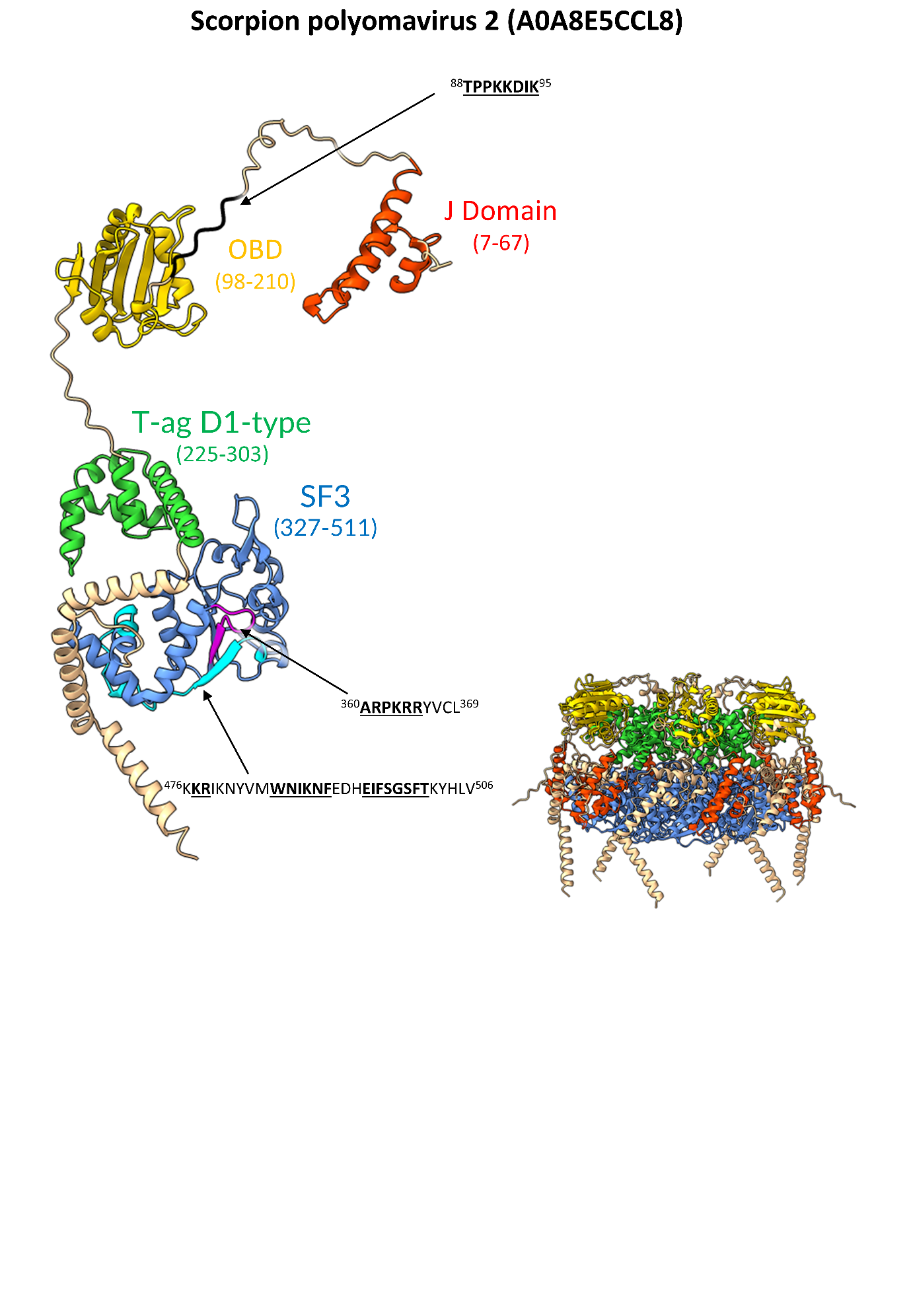
**Supplementary Figure S14. Structural model of Scorpion polyomavirus 2 LTA functional domains.** Monomeric and hexameric forms of scorpion PyV 2 LTA protein (UniProt code: A0A8E5CCL8) predicted using AlphaFold3. The specific domains and motifs are colored according to the following scheme: J domain, red; origin-binding domain (OBD), yellow; T-ag D1 type, green; helicase domain (SF3), blue. Residues encompassing these domains are shown alongside the domain label in parentheses. The identified cNLSs have been indicated with arrows, and colored according to the scheme: residues 360-369 (cNLS within the SF3 domain), magenta; residues 476-506 (cNLS within the SF3 domain), cyan; residues 88-95 (ancestral cNLS upstream of the OBD), black. cNLS sequences are shown with residues in bold and underlined representing flexible regions of the structure whilst those not bold or underlined are identified to be part of the predicted secondary structures.

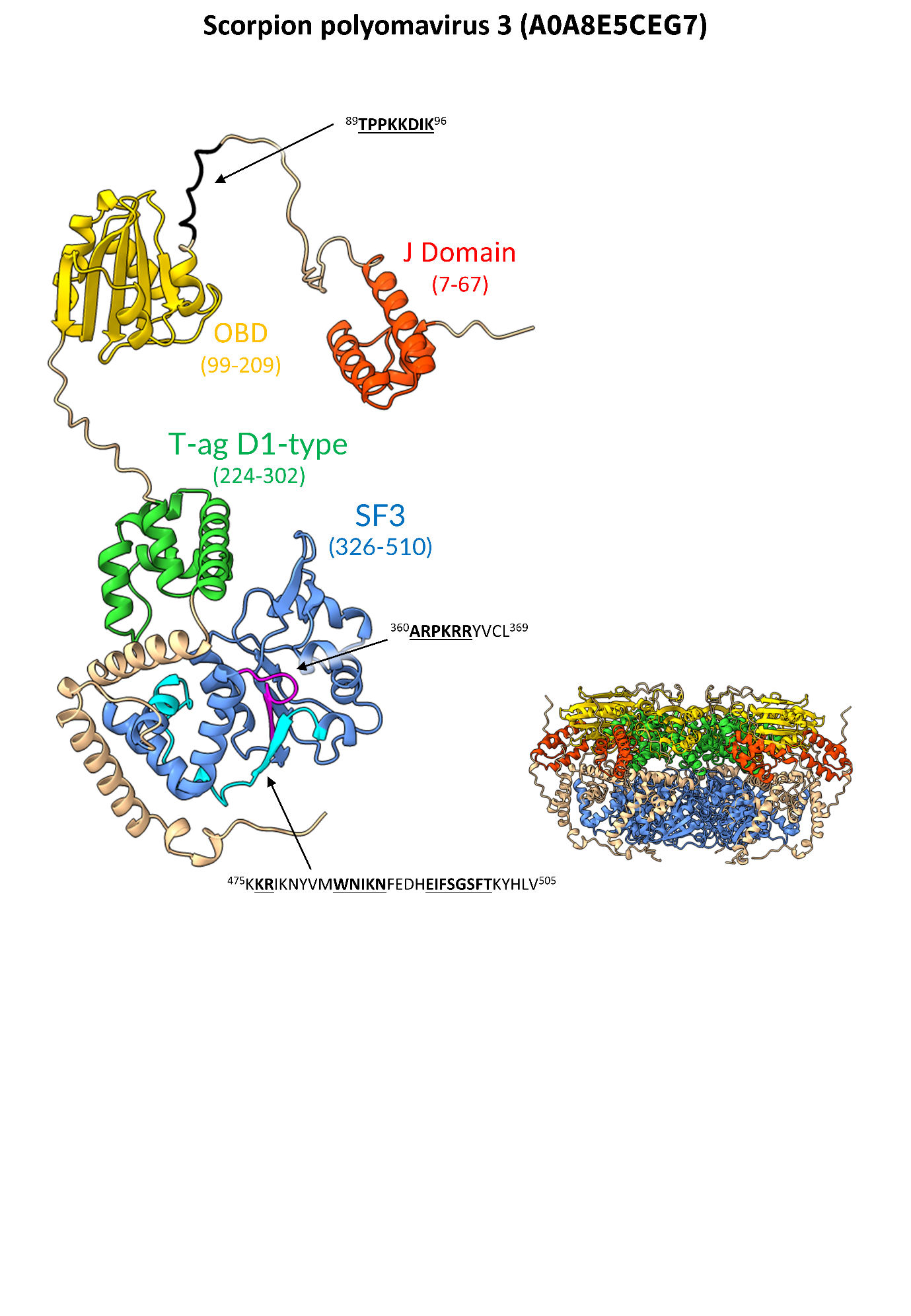

**Supplementary Figure S15. Structural model of Scorpion polyomavirus 3 LTA functional domains.** Monomeric and hexameric forms of scorpion PyV 2 LTA protein (UniProt code: A0A8E5CCL8) predicted using AlphaFold3. The specific domains and motifs are colored according to the following scheme: J domain, red; origin-binding domain (OBD), yellow; T-ag D1 type, green; helicase domain (SF3), blue. Residues encompassing these domains are shown alongside the domain label in parentheses. The identified cNLSs have been indicated with arrows, and colored according to the scheme: residues 360-369 (cNLS within the SF3 domain), magenta; residues 475-505 (cNLS within the SF3 domain), cyan; residues 89-96 (ancestral cNLS upstream of the OBD), black. cNLS sequences are shown with residues in bold and underlined representing flexible regions of the structure whilst those not bold or underlined are identified to be part of the predicted secondary structures.

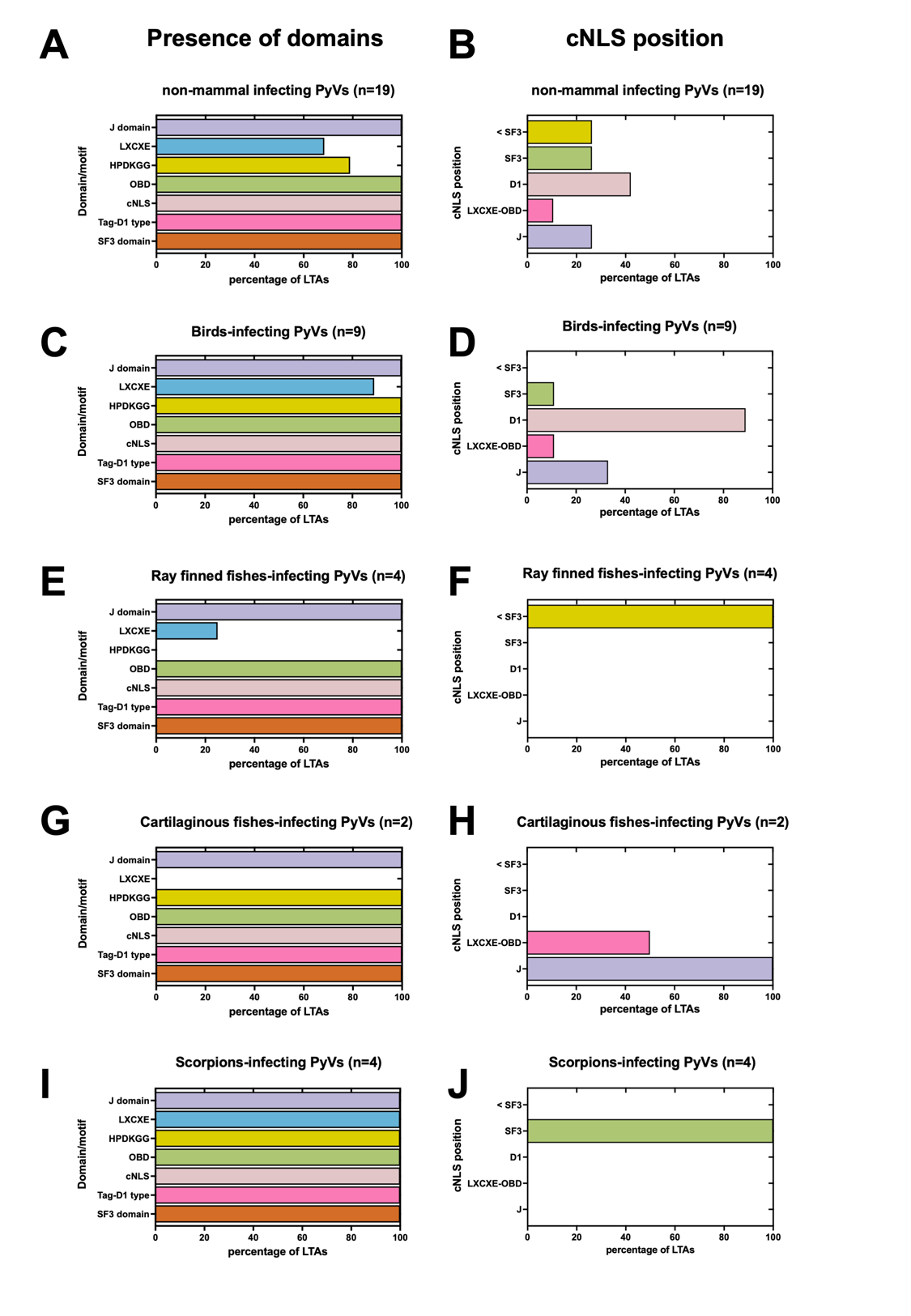

**Supplementary Figure S16 Presence and position of functional domains and motifs in LTAs from PyVs infecting non-mammalian host.** The percentage of LTAs presenting the indicated domain (A, C, E, G, I), and possessing a putative cNLS in the indicated position (D, F, H, J) is shown for all PyVs infecting non mammals (A, B), those infecting birds (C, D), ray finned fishes (E, F), cartilaginous fishes (G, H) and scorpions (I, J). Abbreviations are defined in the Legend of Figure 2.
